# Tom1p ubiquitin ligase structure, interaction with Spt6p, and function in maintaining normal transcript levels and the stability of chromatin in promoters

**DOI:** 10.1101/2024.07.08.601072

**Authors:** Jennifer Madrigal, Heidi L. Schubert, Matthew A. Sdano, Laura McCullough, Zaily Connell, Tim Formosa, Christopher P. Hill

## Abstract

Phosphorylation-dependent binding of the *S. cerevisiae* Spt6p tSH2 domain (Spt6p^tSH2^) to the Rbp1p subunit of RNA polymerase II supports efficient transcription. Here, we report that Spt6p^tSH2^ also binds the HECT-family E3 ubiquitin ligase Tom1p, a homolog of human HUWE1. Tom1p/HUWE1 have been implicated in targeting many small basic proteins for degradation, including excess ribosomal subunits and histones, although the mechanism of substrate recognition is not known. Our cryo-EM data revealed that Tom1p can adopt a compact α-solenoidal “basket” similar to the previously described structure of HUWE1, with the central cavity partially occupied by a disordered acidic domain. Sub-regions of this acidic domain supported binding to Spt6p or histones/nucleosomes *in vitro*, and the histone-binding region was important for Tom1p function *in vivo*. We also visualized Tom1p in more extended forms, and speculate that transitions among these forms could be important for substrate selection and ubiquitylation. Genomic analyses provided additional support for the previously observed role for Tom1p in maintaining ribosomal protein pools, and also demonstrated a role in maintaining chromatin structure near genes. This suggests that the interaction with Spt6p^tSH2^ affects substrate specificity by anchoring Tom1p to localized environments where histone ubiquitylation alters chromatin architecture.

## INTRODUCTION

Spt6p is a multifunctional component of the RNA polymerase II (RNAPII) elongation complex (Vos et al. 2018). A region within the inherently disordered N-terminal domain of Spt6p (Figure 1A) binds to histones, nucleosomes, and the essential transcription factor Spn1p (Diebold et al. 2010, McDonald et al. 2010), the central core domain has been shown to modulate transcription and to contact several elongation complex subunits as well as the emerging transcript (Endoh et al. 2004, Ardehali et al. 2009, Vos et al. 2018, Narain et al. 2021, Zumer et al. 2021), and the C-terminal tandem Src-homology 2 (tSH2) domain assists in the coordination of several transcription-associated activities through direct binding to the Rpb1p subunit of RNAPII (Sdano et al. 2017, Connell et al. 2022) in cooperation with the Cdc73p subunit of the Paf1 Complex (Ellison et al. 2023). The Spt6p tSH2 domain (Spt6p^tSH2^) comprises the only known SH2 motifs in the yeast *Saccharomyces cerevisiae*, and it binds directly to a phosphorylated linker region of Rpb1p between its globular catalytic core and the extended heptad repeats of its C-terminal domain Sdano et al. 2017. This interaction is activated by the Bur1p kinase (Chun et al. 2019), apparently as an early event during the transition between the pre-initiation and elongation phases of transcription, with this interaction contributing to coordination of the dynamic responses of RNAPII to variable local circumstances (Connell et al. 2022).

**Figure 1.**
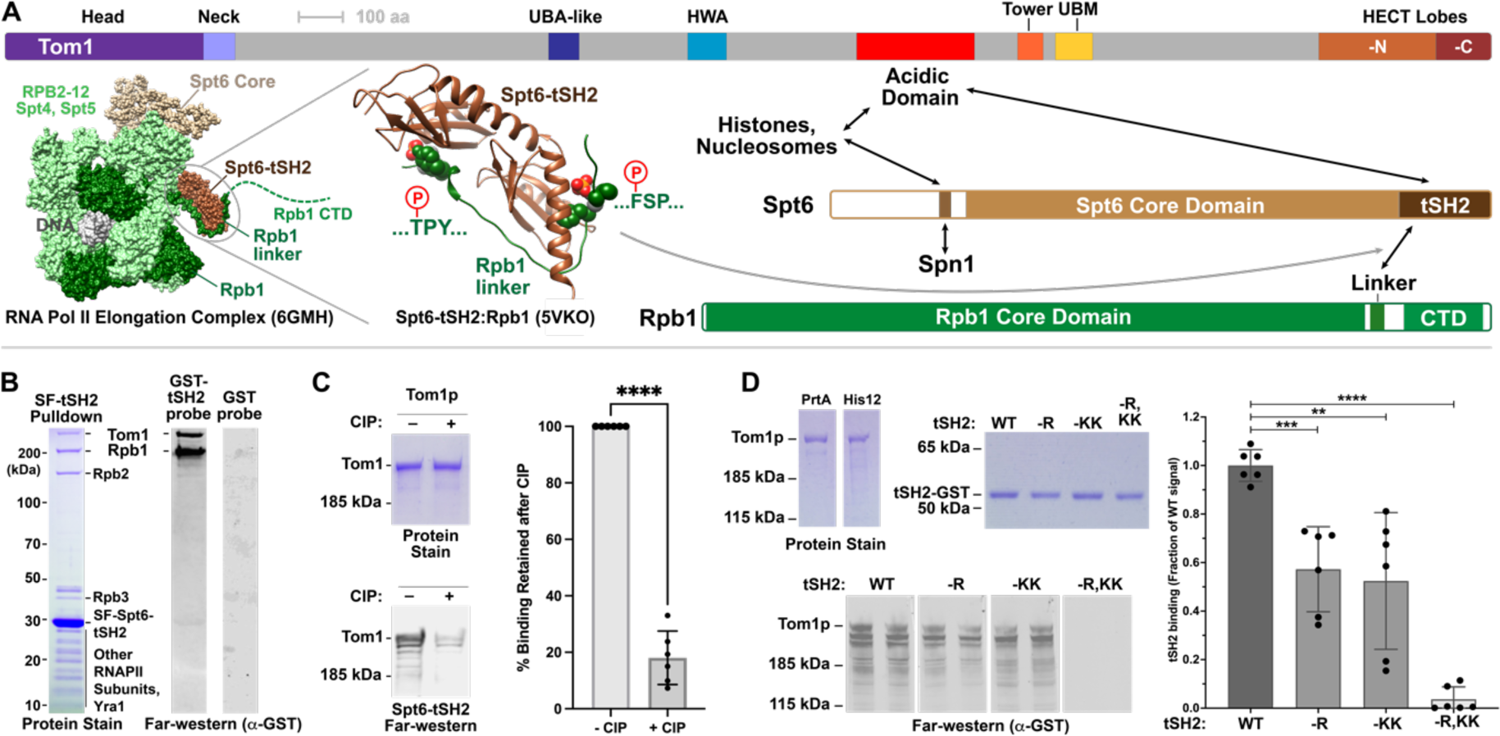
Tom1p is a phosphosphorylation-dependent binding partner of Spt6^tSH2^. **A)** Domain structures of Tom1p, Spt6p, and Rpb1p with 3D structures of the core RNAPII elongation complex (6GMH, Vos et al. 2018) and the tSH2:Rpb1p linker complex (5VKO; Sdano et al. 2017). The two phosphorylation-dependent binding pockets on Spt6p^tSH2^ are shown with phosphates colored red. Arrows indicate interactions of Tom1p, Spt6p and Rpb1p. Domains of Tom1p described in subsequent figures are named. **B)** Yeast strain 7382-3-4 was transformed with vector (Streptavidin-FLAG alone) or plasmid pMS14 (Streptavidin-FLAG-Spt6p^tSH2^), and whole cell lysates were subjected to tandem affinity purification. Proteins that copurified with the Spt6-tSH2 domain were visualized by Coomassie Blue stained SDS-PAGE (Protein) or by detection with purified GST or GST-Spt6p^tSH2^ and anti-GST antibodies (Far-western) after transfer to nitrocellulose. Mass spectroscopy was performed on bands excised from the gel after treatment with trypsin, revealing the identities of the proteins indicated. **C)** Purified Tom1p (Tom1p-Protein A after Protein A removal) was treated with phosphatase (CIP) as indicated. Parallel SDS-PAGE replicas were stained with Coomassie Blue (Protein) or transferred to nitrocellulose and probed with Spt6p^tSH2^ (Far-western). Three repeats were performed with three separate samples, and binding was quantified with each sample normalized to the value obtained without CIP. Overall, phosphatase reduced binding to 18% of the untreated values (P = 4 x 10^-6^). **D)** As in panel C, purified Tom1p was subjected to SDS-PAGE with or without phosphatase (CIP) treatment and probed with WT GST-Spt6p^tSH2^ or variants with mutations in one or both of the binding pockets, see panel A: -R indicates an R1282H mutation in the …TPY… binding pocket, -KK indicates a double K1355A, K1435A mutation in the …FSP… binding pocket, and -R,KK indicates all three mutations as described previously (Sdano et al. 2017). Results from multiple independent measurements were normalized to the average value for WT Spt6-tSH2 with the average and standard deviation shown (t tests yielded P < 0.01 = **, < 0.001 = ***, < 0.0001 = ****).

Herein, we report that in addition to supporting communication among factors associated with RNAPII transcription, the Spt6p^tSH2^ domain also binds directly to the E3 ubiquitin ligase Tom1p. Tom1p is a large, 374 kDa member of a broadly conserved family of HUWE1 factors (also called Mule, Lasu, Ureb1, Arf-BP1, HectH9/E3^Histone^, Ptr1, Upl1, or Eel-1; Hall et al. 2007, Marin 2018). This family is characterized by a large N-terminal domain attached to a smaller C-terminal HECT domain (<u>H</u>omologous to <u>E</u>6-AP <u>C</u>arboxyl <u>T</u>erminus) that provides ubiquitin ligase activity (Figure 1A). Altered HUWE1 function is associated with intellectual disabilities (Froyen et al. 2008, Froyen et al. 2012, Isrie et al. 2013), several forms of cancer (Inoue et al. 2013, Su et al. 2019), dysregulation of responses to general stress and DNA damage (Herold et al. 2008, Thompson et al. 2014, Atsumi et al. 2015, Amici et al. 2022), and aberrant regulation of transcription elongation and transcript processing (Yang et al. 2020, Endres et al. 2021, Solvie et al. 2022).

Tom1p has also been implicated in a broad range of processes in yeast, including maintenance of nuclear organization (Utsugi et al. 1999), cell cycle regulation (Hall et al. 2007, Kim et al. 2012, Kim et al. 2012, Nakatsukasa et al. 2018), ribosomal assembly and quality control (Tabb et al. 2001, Sung et al. 2016, Defenouillere et al. 2017, Pillet et al. 2022), mRNA export (Duncan et al. 2000, Andoh et al. 2004, Iglesias et al. 2010), turnover of free histones (Singh et al. 2009), regulation of transcription (Saleh et al. 1998), and the DNA damage response (Wang et al. 2017). A potential unifying theme for these diverse roles is that many of the substrates of Tom1p/HUWE1 are subunits of large complexes, suggesting a role in turning over unincorporated or defective components (Sung et al. 2016, Xu et al. 2016). Both Tom1p in yeast and HUWE1 in humans have recently been reported to have E4 ubiquitin ligase activity in addition to E3 activity, suggesting a role in enhancing the polyubiquitylation efficacy of other E3 enzymes (Hwang et al. 2010, Ilia et al. 2022, Mengying et al. 2023). It has been proposed that the large N-terminal domain of Tom1p/HUWE1 promotes substrate recognition (Grabarczyk et al. 2021, Hunkeler et al. 2021), but it remains unclear how diverse substrates are recognized by this domain or how the ubiquitylation activity is restricted to appropriate contexts.

In this study, we report finding a direct phosphorylation-dependent physical interaction between Tom1p and Spt6p. Mutational analysis showed that binding requires the same phosphate-binding pockets of Spt6p^tSH2^ that are used to bind the Rpb1p linker. The bound region maps to a conserved acidic domain of Tom1p, a separate portion of which is also important for binding histones and nucleosomes, which are likely to be substrates of Tom1p-mediated ubiquitylation.

Genomic studies supported a pattern of localization of Tom1p that correlates with transcription units and suggested that it has a role in maintaining patterns of nucleosome occupancy that are characteristic of promoters and gene bodies. We determined cryo-EM structures of Tom1p to reveal a helical repeat solenoid architecture for the 2,888 residues preceding the C-terminal HECT domain and an overall architecture that resembles the published structures of HUWE1 but with a larger fraction of ordered residues. In addition to the compact ring/solenoid structure (3.07 Å resolution), we also determined “intermediate” (7.79 Å), and “open” (7.10 Å) conformations that may be important for function. The conserved acidic domain was unstructured, but its ends mapped to an internal face of the solenoid, suggesting that this domain partially occupies the central basket region. An attractive hypothesis is that part of the acidic domain participates in substrate recognition or cofactor recruitment, while surfaces of the solenoid and/or protruding domains also contribute to association with substrate and/or ubiquitin, providing access of substrates to the HECT domain for catalysis.

## RESULTS

### Spt6p^tSH2^ binds directly to Tom1p

We previously used targeted proteolysis, far-western blotting, and crystallography to characterize the interaction between the Spt6p^tSH2^ domain and a linker region in Rpb1p (Figure 1A; Sdano et al. 2017), which subsequently guided analysis and interpretation of RNAPII elongation complex cryo-EM structures (Vos et al. 2018). To ask whether other factors bind Spt6p^tSH2^, we expressed this domain fused to dual streptavidin and FLAG affinity tags in yeast cells and purified it from whole cell lysates under mild conditions. As expected, multiple subunits of the RNAPII complex copurified with the tagged Spt6p^tSH2^ domain, and a far-western blot of these factors probed with a purified Spt6p^tSH2^-GST fusion protein confirmed a direct interaction with Rpb1p (Figure 1B). A single additional factor was also observed to copurify with Spt6p^tSH2^ and to bind the Spt6p^tSH2^-GST probe in the far western blot. Mass spectrometry identified this factor as the 374 kDa Tom1p protein.

To confirm this identification, we replaced the *TOM1* promoter with the inducible *GAL1* promoter at the endogenous genomic locus, fused a Protein A tag at the C-terminus of the reading frame, then used this tag to purify overexpressed Tom1p from yeast cells. Following removal of the tag with PreScission protease, the purified protein (which retained a 14-residue C-terminal scar with the sequence RRIPGLINLEVLFQ) also bound the Spt6p^tSH2^-GST fusion in a far-western assay, thereby validating the results from the affinity copurification experiment (Figure 1C). Binding was substantially reduced upon treatment with phosphatase or when the phosphate-binding pockets in Spt6p^tSH2^ were mutated (Figure 1C, D). The primary direct targets for Spt6p^tSH2^ binding in whole cell lysates therefore appeared to be RNAPII-Rpb1p and Tom1p, and these interactions had similar requirements for phosphorylation of the binding sites on Rpb1p or Tom1p and the integrity of the Spt6p^tSH2^ pockets that bind Rpb1p peptide phosphoryl groups (Sdano et al. 2017).

### The conserved central acidic domain of Tom1p contains the Spt6p^tSH2^ binding site

Partial proteolysis of purified Tom1p with trypsin coupled with far-western blotting localized the Spt6p^tSH2^ binding site to an ∼60 kDa trypsin-resistant fragment (Figure 2A). The majority of unique peptides identified from this band by mass spectrometry clustered in a region adjacent to the large, conserved acidic domain found in all members of the HUWE1 family, which would be expected to be resistant to trypsin (Figure 2B). We therefore deleted the 259-residue acidic domain (Tom1p^ΔAD^; lacking E1873-E2131; 38% D, E, 4% K, R, pI 3.0) and found that the resulting protein no longer bound Spt6p^tSH2^ in the far-western assay (Figure 2C). Tom1p^ΔAD^ was expressed at the normal level in yeast and the purified protein was well-behaved (Figure 2C), so deletion of the acidic domain did not destabilize Tom1p but did substantially reduce the affinity for Spt6p^tSH2^. Additional internal deletions of segments of the acidic domain were constructed and tested, localizing the Spt6p^tSH2^ binding site to a region of 33 amino acids (*tom1-intΔ11* lacking 2031-2063, Figure 2C).

**Figure 2.**
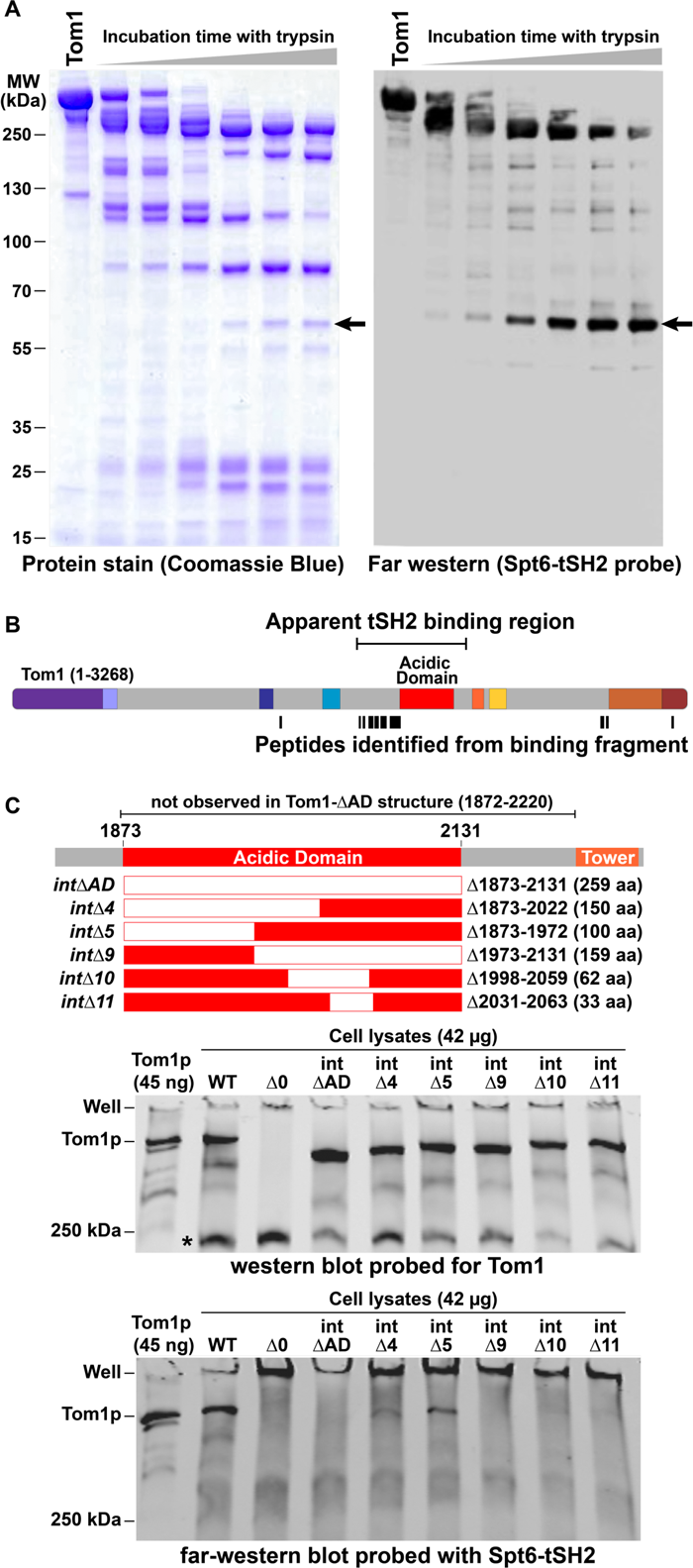
Spt6p^tSH2^ binding maps to the acidic domain of Tom1p. **A)** Purified Tom1p was incubated with trypsin. Aliquots removed at the indicated times were visualized by SDS-PAGE followed by Coomassie Blue staining (left) or far-western probing with GST-Spt6p^tSH2^. **B)** The 60 kD band indicated with an arrow in (A) was excised and subjected to complete digestion with trypsin followed by mass spectrometry. The locations of peptides identified are shown in panel B. The inferred tSH2 binding site (bracket) starts with the first peptide of the most significant cluster of peptides and extends about 60 kDa, including the acidic domain which contains few cleavage sites for trypsin. **C)** Markerless in-frame deletions were constructed to remove the entire acidic domain (*intΔAD*) or subregions within this domain at the endogenous *TOM1* locus. Whole cell lysates were prepared from log-phase cultures and subjected to SDS-PAGE followed by detection with antibodies against Tom1p (middle panel) or Spt6p^tSH2^-GST fusion protein (bottom panel). The antisera cross-reacted with an unidentified protein of about 250 kDa (asterisk) which was unrelated to Tom1p as it was retained in a strain lacking the entire *TOM1* gene (*tom1-Δ0*).

### The acidic domain comprises separable regions required for individual Tom1p functions

Deletion of the complete Tom1p reading frame (the *tom1-Δ0* allele) causes tight temperature sensitivity (Utsugi et al. 1999 and Figure 3). This might result from a general defect in processing misfolded proteins that accumulate at higher temperatures, although some observations challenge this interpretation, including the abrupt onset of the growth defect (growth was normal at 30°C, weak at 33°C, and blocked at 34°C; Figure 3) and partial suppression of the phenotype by use of a different carbon source (*tom1-Δ0* failed to grow on glucose at 35°C but was less impaired when grown on galactose at the same temperature; Figure 3). Other phenotypes caused by complete loss of Tom1p included sensitivity to caffeine, formamide, and the DNA-damaging agent phleomycin (Figure 3). Notably, deletion of just the acidic domain (*tom1-intΔAD*) or the left half of this region (*tom1-intΔ4*) caused a similar spectrum of phenotypes as the full gene deletion, even though these alleles did not strongly alter the level of Tom1p in lysates (Figure 2C). The N-terminal portion of the acidic domain therefore appears to be essential for the *in vivo* functions of Tom1p that are needed to prevent these phenotypes. Deleting less of this region in *tom1-intΔ5* caused a more moderate growth defect at 34°C (although a more severe defect at higher temperatures), and deletions that removed the right side of the acidic domain displayed no apparent phenotypes in these plate assays. The acidic domain therefore appears to have specific functional regions, parts of the left side of the region may have additive or modular roles, and the loss of the site responsible for the main interaction with Spt6p^tSH2^ (*tom1-intΔ10, 11*) did not result in failure of a Tom1p function detectable by these growth assays.

**Figure 3.**
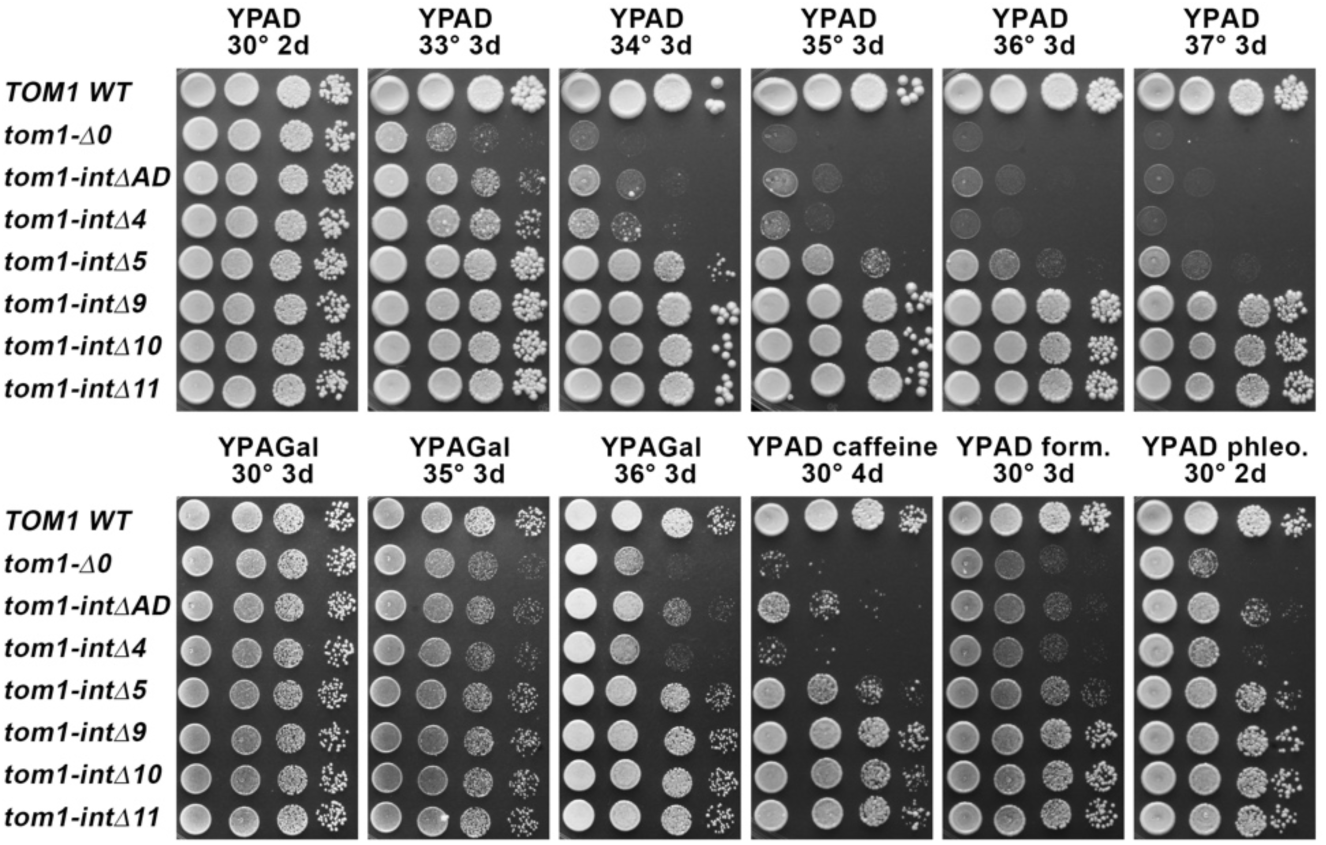
The Tom1p acidic domain is important for *in vivo* functions. Yeast strains with the mutations shown in Figure 2C were grown to saturation, washed in water, and 10-fold serial dilutions were placed on the media shown and incubated as indicated. YPAD is yeast extract and peptone supplemented with adenine and 2% dextrose and, as indicated, with 15 mM caffeine, 3% formamide, or 6 µg/mL phleomycin. Gal indicates substitution of the dextrose with galactose.

### Tom1p binds histones and nucleosomes through the acidic domain

Tom1p has been implicated in the turnover of excess levels of small, basic proteins including ribosomal components (Tabb et al. 2001, Sung et al. 2016) and histones (Singh et al. 2009). Unincorporated individual ribosomal proteins are generally unsuitable for binding assays due to their insolubility, so to test the importance of the acidic domain of Tom1p for binding potential substrates we used EMSA with soluble histone complexes. As shown in Figure 4, Tom1p formed complexes with H2A-H2B dimers, (H3-H4)_2_ tetramers, and also with assembled nucleosomes. Titrations of Tom1p or H2A-H2B gave K_d_ values of ∼160-330 nM in this assay (Figure S4). Binding was not observed with Tom1p^ΔAD^ but complexes formed normally when just the Spt6p^tSH2^ binding site was removed in Tom1p^Δ11^ (Figure 4). Taken together, the results support the conclusion that the acidic domain is important for binding potential substrates like histones, that this is important for Tom1p function *in vivo*, and that the interaction with Spt6p^tSH2^ has a role distinct from general substrate recognition.

**Figure 4.**
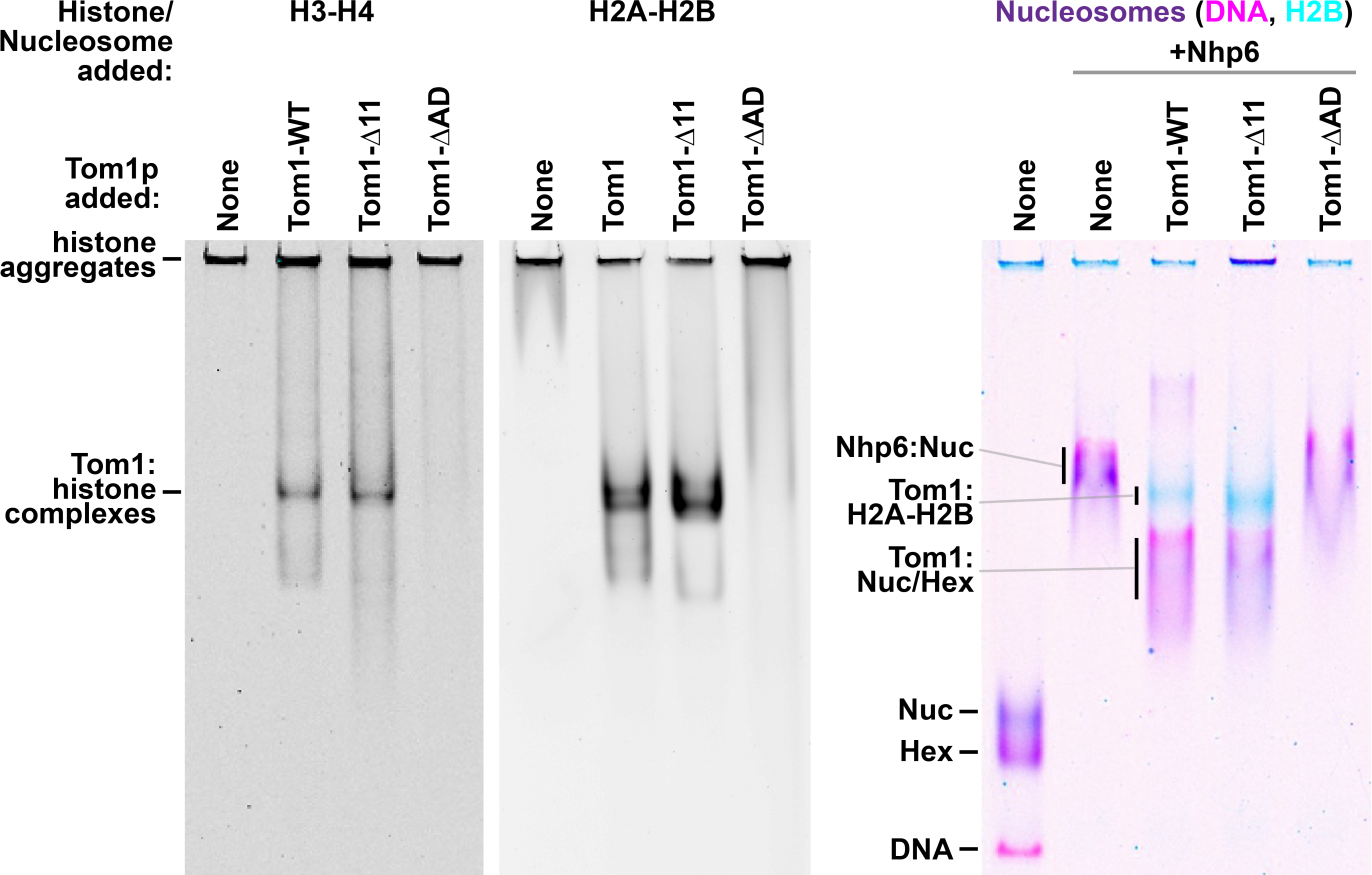
The acidic domain of Tom1p mediates binding to histones and nucleosomes. Versions of Tom1p fused to Protein A were purified as described in Methods, mixed with purified yeast histones alone or after reconstitution into nucleosomes with a 181 bp DNA fragment, then separated by native PAGE (Xin et al. 2009). Histones and nucleosomes were detected by a fluorescent dye attached to a unique cysteine residue in H3 or H2B, or at the 5’ end of the DNA. Free histones do not migrate uniformly in this gel system, typically forming aggregates in the well or being lost during analysis, so bands indicate the formation of complexes with Tom1p. Nucleosomes retain variable amounts of H2A-H2B dimer, with octamers (nucleosomes) and hexamers (hexasomes) migrating differentially, as indicated. Similar to the results with FACT and Spt6p (Xin et al. 2009, McCullough et al. 2015), nucleosomes only formed complexes with Tom1p in the presence of the HMGB family factor Nhp6p.

### Genomic analysis supports a role for Tom1p in managing chromatin during transcription

Interaction of Tom1p with Spt6p and histones suggests a role in transcription, possibly assisting Spt6p with managing the properties of chromatin as it is dynamically remodeled during promoter activation or transcription elongation. To test this possibility, we measured the effects of deleting *TOM1* on steady-state transcript levels by RNA-seq, the global occupancy of Tom1p by ChIP-seq and the effects of deleting *TOM1* on chromatin structure by MNase-seq.

RNA-seq revealed a significant decrease in the steady-state level of ribosomal protein gene transcripts in cells lacking Tom1p (Figure 5A). This is consistent with a role for Tom1p in degrading unincorporated ribosomal subunits as previously described (Sung et al. 2016) as unbalanced expression of any ribosomal protein signals a reduction in the transcription of all genes in this class.

**Figure 5.**
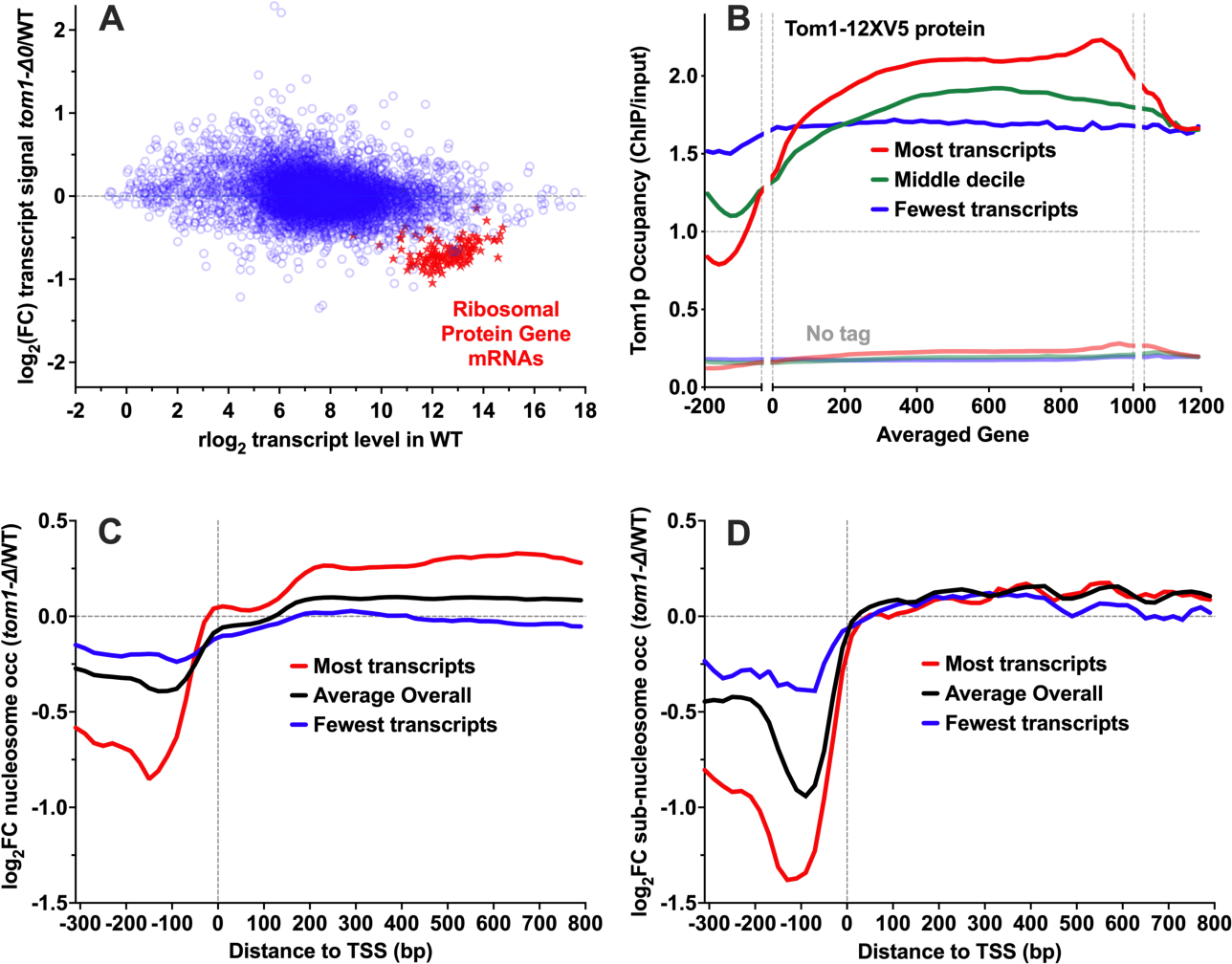
Genomic analysis supports a role for Tom1p in transcription that involves maintaining chromatin architecture in promoters and gene bodies. **A)** RNA-seq was performed with WT and matched *tom1-Δ0* strains in triplicate and the log2FC value for mutant/WT was calculated for each gene. Transcript levels from most genes were only modestly affected by loss of Tom1p (7 genes had decreased transcript levels and 16 genes were increased at a threshold of a 2-fold decrease at a false discovery rate below 1%). As a group, transcripts from genes encoding ribosomal protein genes (red stars) were generally decreased, forming a distinct subset (red stars). **B)** A 12xV5 tag was fused to the C-terminus of the Tom1p ORF at the endogenous locus and anti-V5 antibodies were used to perform ChIP-seq with three independent clones and a matched strain without the tag. ChIP signal relative to input DNA was calculated genome-wide and mapped for each gene normalized to an averaged gene size of 1000 arbitrary units, as well as 200 bp upstream and downstream of each gene. Genes were ordered by their transcript levels determined by RNA-seq using a matched WT strain (Connell et al. 2022 and panel A). The average Tom1p occupancy values are shown for the highest, lowest, and middle transcript level deciles. **C, D)** MNase-seq was performed in triplicate with independent isolates of WT and *tom1-Δ0* strains, and reads were parsed into groups by the size of the protected DNA fragment. 140-170 bp fragments were classified as nucleosomes, representing ∼36% of all reads, and 90-120 bp fragments were classified as subnucleosomes, representing ∼3.6% of all reads. Protected fragments were mapped to the annotated transcription start site (TSS) of each gene. Genes were placed in deciles by transcript levels as in panel B and the log_2_FC occupancy values shown are relative to nucleosomes (C) and subnucleosomes (D).

Antisera generated against purified Tom1p were not suitable for ChIP, presumably due to interference from an abundant cross-reacting protein (asterisk, Figure 2C). Epitope tags fused to either the N-terminus or C-terminus of Tom1p produced the same spectrum of phenotypes as null alleles, suggesting inactivation of key functions of Tom1p (not shown). We therefore used a 12x-V5 tag fused to the C-terminus of Tom1p for the ChIP-seq experiment; while this disrupted its physiological function (consistent with results obtained with other HECT domain proteins like E6AP and Rsp5p; Salvat et al. 2004, Kamadurai et al. 2009), it did not alter the steady-state level of Tom1p or its interaction with Spt6, suggesting any Spt6-driven localization should remain intact.

Tom1p-12xV5 occupancy mapped to gene bodies, especially genes with high transcript levels (Figure 5B). In contrast, occupancy was particularly low over the promoters of these highly expressed genes. An untagged strain gave little signal, so we conclude that Tom1p was associated with highly transcribed gene bodies but was less associated with promoters.

To examine a potential role in maintaining chromatin, we mapped nucleosomal occupancy by paired-end MNase-seq in normal and *tom1-Δ0* strains (Figure 5C). Once again, we observed stronger effects in the decile of genes with the highest transcript levels, with loss of Tom1p leading to a decrease in nucleosome occupancy relative to a WT strain over promoters, approximately normal occupancy at the +1 nucleosome, and elevated occupancy over gene bodies. This suggests that Tom1p has a role in stabilizing nucleosomal occupancy in promoters and slightly decreasing the occupancy in gene bodies. A similar effect was seen over promoters when we mapped MNase-resistant DNA fragments smaller than those protected by full nucleosomes (“sub-nucleosomal” fragments that could arise from partial nucleosomes or other DNA-binding proteins) (Figure 5D). In this case, gene bodies were relatively unaffected, but loss of Tom1p caused a universal reduction in protection of promoter regions. Tom1p therefore appears to have a role in maintaining chromatin architecture, especially in the promoters of highly transcribed genes.

*TOM1* has been implicated previously in histone turnover (Singh et al. 2009), but we did not observe a change in the total steady-state level of either soluble or chromatin-bound histone H3 in strains lacking Tom1p (Figure S5). A role in histone turnover may therefore be redundant with other factors or limited to a specific subset of histones, possibly those being displaced near promoters as the result of dynamic processes related to transcriptional regulation.

Collectively, these genomics results are consistent with a role for Tom1p in maintaining normal levels of ribosomal proteins and transcripts. They also show that Tom1p occupancy varies with gene structure, especially for highly transcribed genes, and it may directly affect the stability or turnover of nucleosomes, with differential effects in promoters and gene bodies.

### Tom1p Structure Determination

We performed single-particle cryo-EM reconstructions of two Tom1p constructs: full-length Tom1p that contained a 20-residue tag including 12 histidines inserted between residues K1074-D1075 (Tom1p^His^; physiologically active in all phenotype assays) and Tom1p^ΔAD^ with the C-terminal Protein A described for intact Tom1p above (Figure 2C, presumably inactive). These structures were determined at resolutions of 3.97 Å (Tom1p^His^) and 3.07 Å (Tom1p^ΔAD^) for the dominant closed ring conformations (Figure 6B, 7C, S6). The two maps and structural models were very similar to each other, indicating that neither the internal affinity tag, the absence of the acidic domain, or the presence of a C-terminal Protein A scar induced notable conformational changes. Unless otherwise noted, the higher resolution Tom1p^ΔAD^ map was used for descriptions and illustrations herein. As expected, the acidic domain was disordered in the Tom1p^His^ structure. 2,792 (85%) of the 3,268 Tom1p residues were modeled into Tom1p^ΔAD^, which corresponds to 96% of the residues in Tom1p^ΔAD^ (Figure 6A).

**Figure 6.**
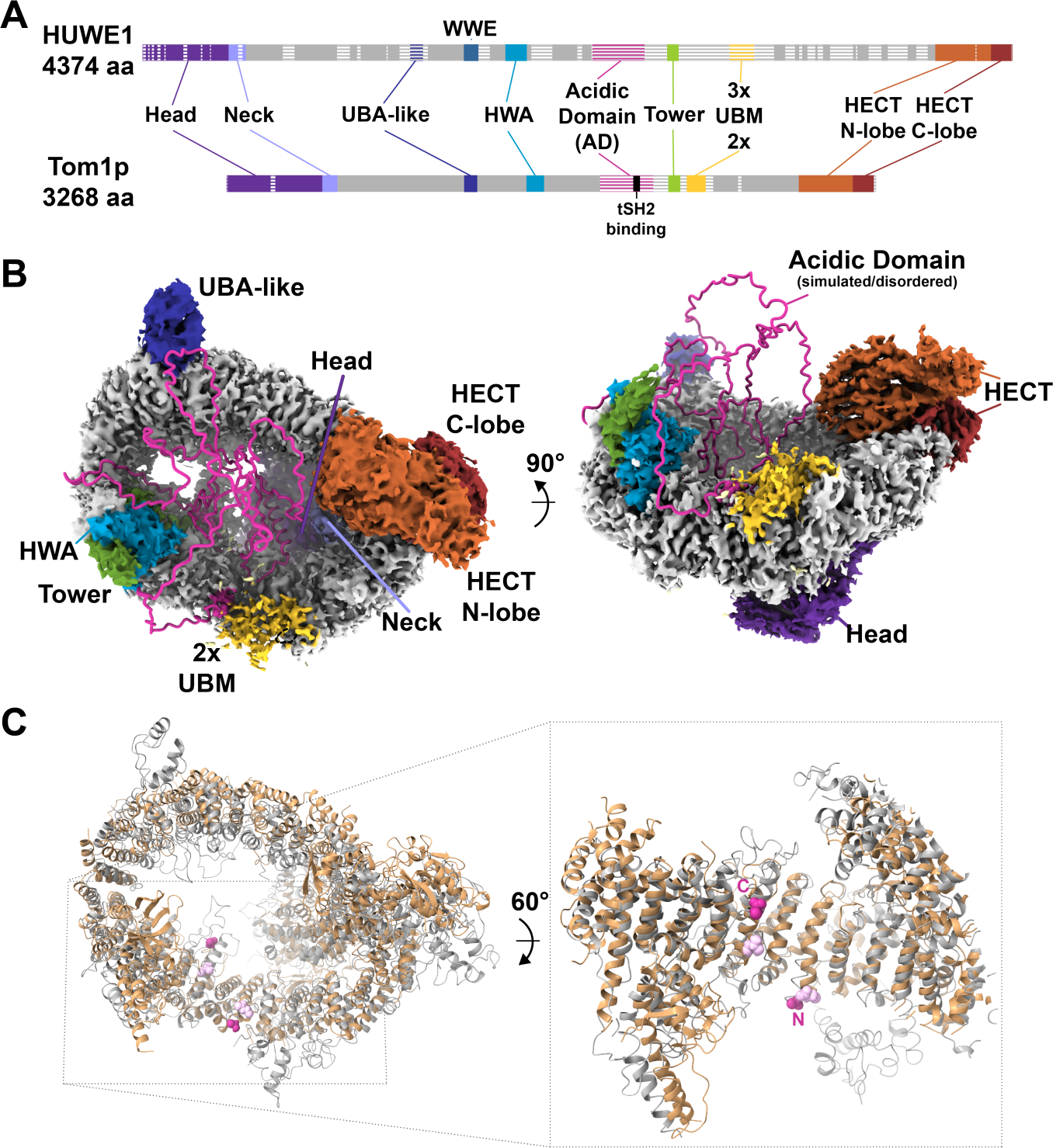
Structure of yeast Tom1p in the closed conformation and comparison with human HUWE1. **A)** Alignment of human HUWE1 and Tom1p sequences. Modeled residues indicated with solid bars; disordered regions are striped. Motifs/domains recognized from the sequence or defined in the structure are indicated with color. Tom1p: Head domain (1-436), Neck (437-506), UBA-like domain (1195-1261), HWA(1510-1594), acidic domain (1873-2131), Tower helix (2221-2276), UBM module (2317-2398), HECT N-terminal lobe (2889-3146), HECT C-terminal lobe (3147-3268). Body colored grey. Human HUWE1 WWE domain is not present in Tom1p. Beginning and end points of missing density in Tom1p^ΔAD^ structure: 214-235, 1873-2219 (region containing acidic domain), 2276-2316, 2399-2449, 2575-2589. **B)** Top-down (left) and side (right) views of the 3.07 Å Tom1p^ΔAD^ map. Disordered residues of the acid domain are shown as pink coil in random positions. Other disordered regions not depicted. The structure of Tom1p^His^ is very similar, with the exception of more poorly resolved density at the UBM domain. **C)** Top-down view of Tom1pΔAD (grey) aligned with human HUWE1 (tan) (Hunkeler et al. 2001; PDB 7JQ9). The close-up shows a clipped side view of the interior of the basket between the Tower and UBM regions. The ordered residues either side of the disordered acidic domain are shown: Tom1p, magenta, residues 1872, 2220; HUWE1 light pink).

### Closed Ring Conformation and Comparison with HUWE1

Similar to the published structures of HUWE1 from humans (Hunkeler et al. 2021) and the fungus *Nematocida* (Grabarczyk et al. 2021), Tom1p comprises more than 100 helices in a series of HEAT or ARM-like repeats that extend over the first 2,888 residues in an α-solenoid conformation comprised of Head, Neck, Body that precedes the C-terminal HECT domains (Figure 6).

The N-terminal 436 residues comprise the Head domain, in which the first 6 helices form three 2-helix repeats, followed by a bundle of loops and helices from residues 153-286, before the helical repeat pattern continues at residue 287 with several ARM repeats. The globular Head is followed by the Neck domain (residues 437-506) comprising one ARM-like repeat followed by a loop (residues 488-506) that extends toward the interior of the solenoid to lie along the surface of the Head domain (Figure S6C). The Neck is followed by the Body domain (residues 507-2,888), which comprises a helical repeat solenoid structure that forms a ring architecture and leads into the C-terminal HECT domain. The first helix of the HECT domain, residues 2889-2906, helps to close the ring by packing between the HECT N- and C-lobes on one side and the Head and Neck domains on the other side. The Head domain largely closes one side of the ring formed by the Body domain to create a basket-shaped structure, while the HECT domain is positioned to one side at the opposite, open face of the basket. This overall structure is conserved in human HUWE1 (Figure 6A, 6C), although one-to-one structural correspondence of individual residues is not generally apparent outside of the HECT domain. This is consistent with the low (20%) sequence identity for *S. cerevisiae* Tom1p and human HUWE1 residues outside of the HECT domain and higher (53%) identity between their HECT domains.

Several domains, including some with similarity to known protein-interaction modules, are encoded in inter-helical loops that protrude from the Body domain on the rim of the basket structure. Some of these motifs and domains have been identified in the sequence of human HUWE1, although several of them were not visualized in the HUWE1 structure due to disorder (Grabarczyk et al. 2021). Most of the 1,106 additional residues in the 4,373-residue HUWE1 protein that lack counterparts in Tom1p are disordered in the HUWE1 structure and occur, in about equal parts, as insertions between the solenoid helices or within the protrusions that are conserved in Tom1p.

All of the protruding domains in Tom1p have counterparts in HUWE1. Starting from the N-terminal end, these include a 3-helix bundle (residues 1201-1260) that corresponds to a region of HUWE1 that was classified as a UBA (Ubiquitin Associated) domain on the basis of sequence alignment, although these residues were not visualized in the HUWE1 structure and in Tom1p the third helix adopts the opposite orientation relative to canonical UBA domains. An HWA (HUWE1 Associated) domain is conserved in HUWE1 and Tom1p (residues 1510-1594), although the structurally adjacent 78-residue WWE domain of HUWE1 is replaced by a loop in Tom1p (residues 1404-1427). The Tom1p acidic domain (residues 1873-2131), which is conserved in HUWE1, is found within a larger disordered region (residues 1871-2219), the ends of which are located on the inner face of the solenoid, suggesting that this unstructured component of Tom1p will at least partially occupy the basket (Figure 6B, 6C). Ordered structure resumes only six residues before the start of the Tower domain (residues 2221-2276), for which the two-helix protrusion of HUWE1 is replaced by a single helical protrusion in Tom1p followed by ∼50 flexible residues and 2 adjacent UBMs (Ubiquitin-Binding Motifs; residues 2317-2398) that substitute for the 3 UBMs observed in HUWE1 (Figure 6A).

### Open conformation

In addition to the closed ring structure, the Tom1p^His^ dataset contains particles in “intermediate” and “open” conformations that were reconstructed to resolutions of 7.79 and 7.10 Å, respectively (Figs 7C, S6C-D). The distinction between closed and open/intermediate conformations is modeled as a rigid body hinging of 30° (open) or 15° (intermediate) centered at a loop (1316-1320) in the middle of the Body domain at the opposite side of the ring from the N-terminal Head and C-terminal HECT domains (Figure 7B, 7C). The resolution does not allow precise modeling of the change in conformation, but apart from the rigid body rotations, conformational changes appear to be limited to some repacking of the interface between the two halves of the Body domain and modest shift in adjacent helices (Figure 7A).

**Figure 7.**
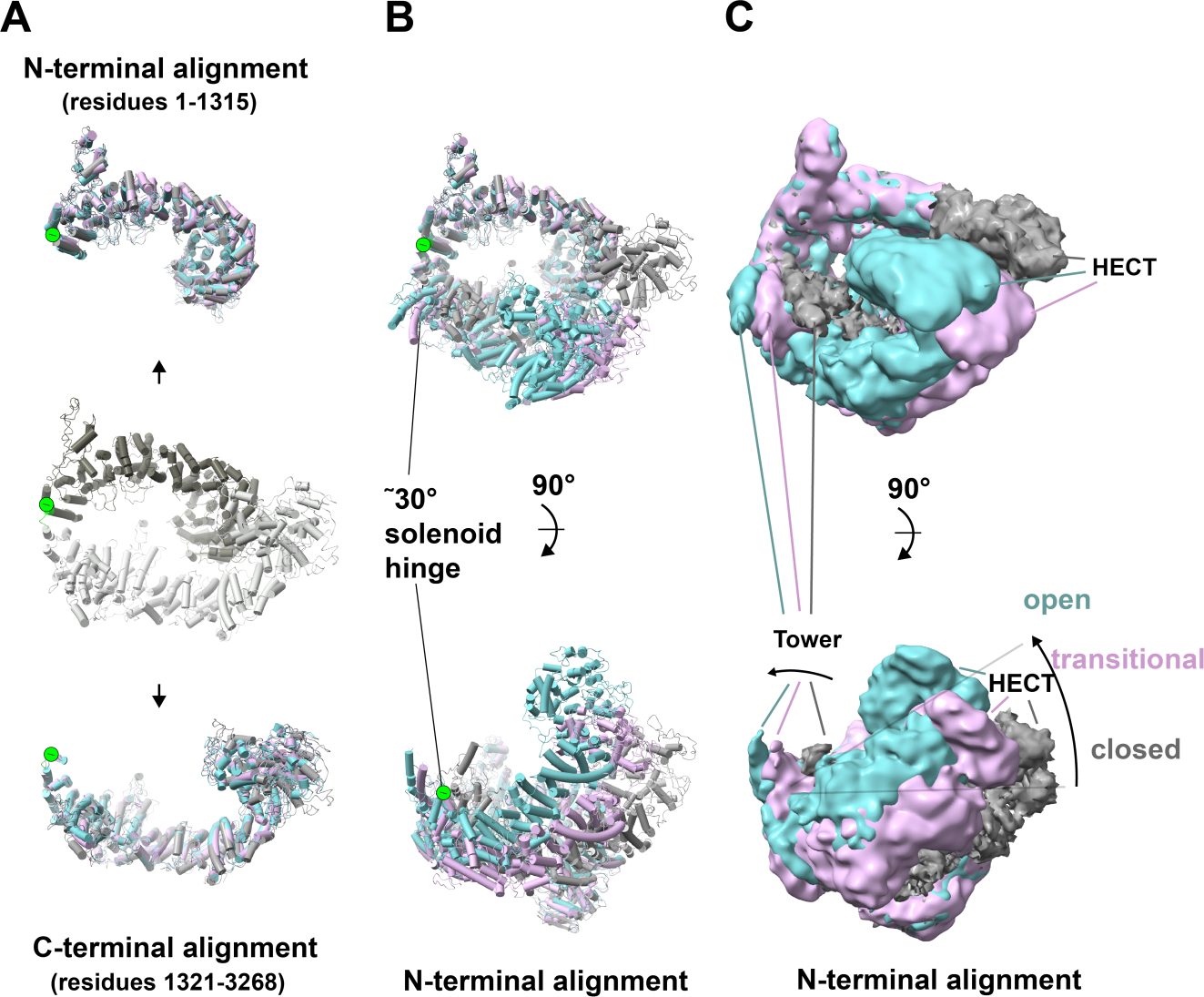
Tom1p^His^ hinging motion. **A)** When aligned separately, the N-terminal and C-terminal halves of closed, intermediate, and open conformation Tom1 superimpose closely. Center: Top-down view of the Tom1p^His^ closed-conformation model with residues 1-1315 colored dark grey and 1321-3268 colored light gray. Top: same orientation showing the three Tom1p^His^ conformation models aligned for residues 1-1315: closed (grey), intermediate, (pink) and opened (turquoise). Bottom: Same orientation, showing the three Tom1p^His^ conformation models aligned for residues 1321-3268. **B)** The N-terminal and C-terminal halves of closed, intermediate, and open conformation Tom1p display a hinging motion Top-down and side views of the three Tom1p^His^ conformation models aligned on residues 1-1315. The hinging motion, centered on loop 1316-1320 (green), is ^∽^ 15° and ^∽^30° for the intermediate and open conformations, respectively. **C)** Same as (B) but showing maps for the three structures.

Published cryo-EM studies of human HUWE1 indicated the presence of movement towards an open state analogous to that of Tom1p, although density for residues C-terminal to the hinge were not apparent (Hunkeler et al. 2021). Mutation of two residues in the Head domain at the solenoid closure interface reduced human HUWE1 E3-ligase activity, suggesting that the closed conformation is important for the catalytic cycle (Hunkeler et al. 2021).

### The HECT domain adopts a constrained/inactive conformation

The 379-residue HECT domain adopts similar structures and maintains the same intramolecular contacts in the open and closed solenoid structures of Tom1p and in the published structures of human (Hunkeler et al. 2021) and *Nematocida* (Grabarczyk et al. 2021) HUWE1. Overlap of the Tom1p and human HUWE1 HECT domains shows a 1.1 Å RMSD over 256 pairs of Cα atoms. HECT domains comprise two lobes, an N-terminal lobe and a smaller C-terminal lobe that houses the catalytic cysteine residue (C3235 in Tom1p). Multiple HECT domain crystal structures have indicated that the two lobes rotate with respect to each other through the catalytic cycle of binding E2, transthiolation of ubiquitin from E2 to the HECT domain cysteine, and subsequent ubiquitylation of substrate. The N-terminal Head domain makes clearly defined, extensive contacts with the final ARM-like repeat of the Body domain that leads into the first helix of the HECT domain (residues 2289-2904; Kane et al. 2022). The HECT domain conformation appears to be unaltered in the open or intermediate structures, and we were unable to show substantial HECT domain C-lobe motion relative to the N-lobe, or motion of the HECT domain relative to the neighboring segment of solenoid using 3DVA (Punjani et al. 2021) or the beta version of 3D Flex (Punjani et al. 2023).

The Tom1p HECT conformation closely resembles that of the human apo HECT domains of HUWE1 (PDB ID 3H1D; Pandya et al. 2010), AREL1 (PDB ID 6JX5; Singh et al. 2019), and WWP1 (PDB ID 1ND7; Verdecia et al. 2003), with the HECT catalytic cysteine inaccessible to a bound ubiquitin-charged E2 enzyme (Figure 8A, 8B, 8C). Simple docking of Tom1p onto a NEDD4L HECT domain complex with a ubiquitin-charged E2; Kamadurai et al. 2009, PDB ID 3JW0) or onto HECT-Ub structure (HUWE1:Ubq; Nair et al. 2021, PDB ID 6XZ1) suggests that the Tom1p HECT could undergo a full catalytic cycle within the framework of the closed conformation without major structural adjustment of the N-lobe with respect to the α-solenoid (Figure 8C, 8D).

**Figure 8.**
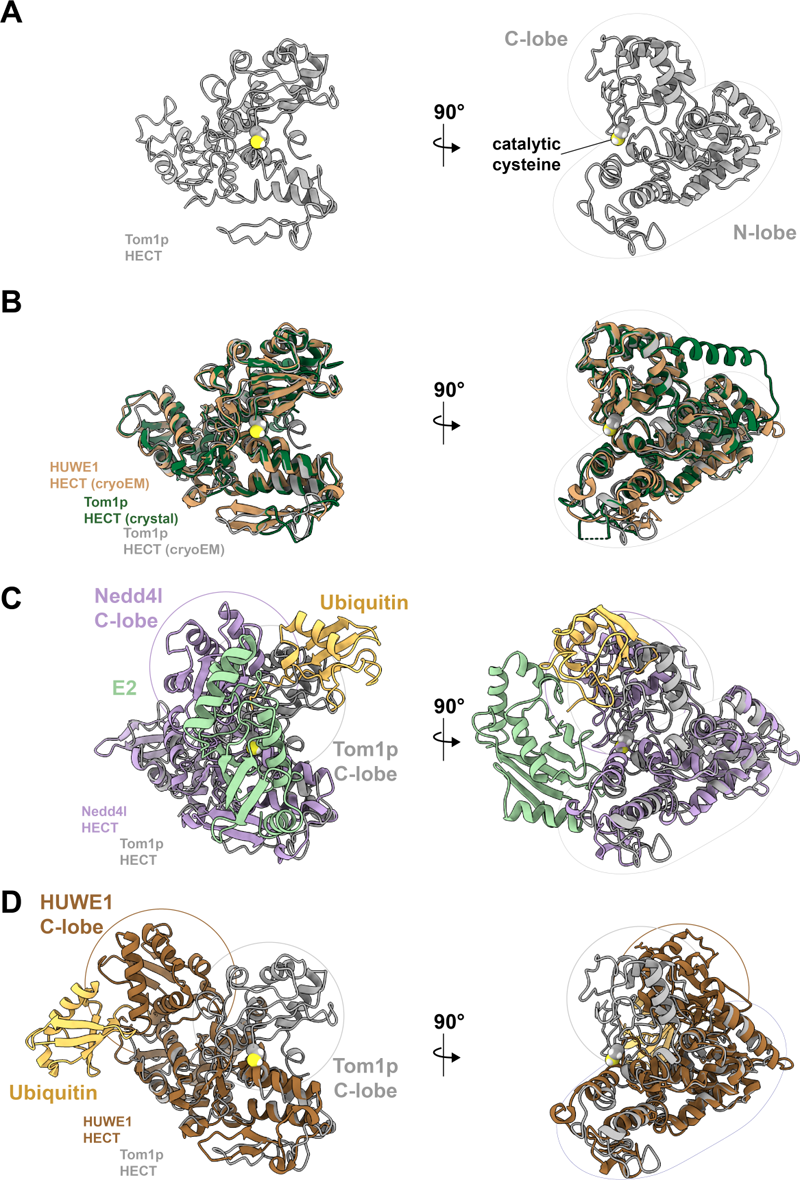
HECT domain comparisons between constructs, conformations, homologs, and catalytic states. **A)** Tom1 HECT domain. Tom1p^ΔAD^ HECT domain is shown. Tom1p^His^ closed, intermediate, and open conformations are superimposable within experimental error. Catalytic cysteine, yellow. **B)** Superposition of Tom1p and HUWE1 HECT domains. Tom1p^ΔAD^, gray; human HUWE1, tan (Hunkeler et al. 2021; PDB 7MWD), Tom1p HECT crystal structure, green. **C)** Tom1p (grey) and human Nedd4L HECT domains superimposed on their N-lobes. The Nedd4L HECT structure (lavender) is a complex with a Ub-charged E2 (yellow, green) (PDB 3JW0; Kamadurai et al. 2009. C-lobes show a relative rotation of 63°. **D)** Tom1p^ΔAD^ (grey) superimposed on N-lobe with human HUWE1 HECT domain (brown) covalently bound to Ub-propargylamine (yellow) (PDB 6XZ1; Nair et al. 2021). C-lobes show a relative rotation of 167°.

## DISCUSSION

Spt6p contacts several subunits of the core RNAPII elongation complex, the emerging transcript, and the surfaces of histones as nucleosomes are transiently disrupted during transcription (Vos et al. 2018, Zumer et al. 2021). The C-terminal tSH2 domain of Spt6p binds a phosphorylated linker region of the RNAPII catalytic subunit, Rpb1p, and this contact is important for coordinating the dynamic changes that occur during a transcription cycle, potentially by mediating communication among the multiple functional modules of the RNAPII elongation complex (Sdano et al. 2017, Connell et al. 2022). Here, we report an unexpected additional Spt6p^tSH2^ ligand, the E3 ubiquitin ligase Tom1p. Similar to the Spt6p^tSH2^-Rpb1p interaction, Tom1p binding was phosphorylation-dependent and mapped to a short peptide within an unstructured region, in this case an acidic domain conserved among known Tom1p/HUWE1 homologs. Genomic analysis supported the previously described role for Tom1p in regulating ribosomal protein turnover but also suggested an unexpected role in transcriptional regulation, possibly through altered histone turnover that affects local chromatin architecture. Our structural analysis revealed a closed solenoidal ring structure and multiple open conformations suggestive of dynamic conformational changes to a spiral shape. These findings provide a foundation for efforts to understand how this conserved family of ubiquitin ligases chooses substrates, how its activity is limited to appropriate circumstances, and how it might modulate chromatin structures important for transcriptional regulation or DNA repair.

The HECT domains of Tom1p/HUWE1 family members are thought to ubquitylate substrates, likely for subsequent degradation by the proteasome, while more N-terminal sequences presumably function in substrate recognition. Our data support a role for the flexible acidic domain in binding small basic proteins like ribosomal proteins and histones, although a simple electrostatic mechanism seems inadequate to produce appropriate specificity. We confirmed that the acidic domain of Tom1p was necessary for binding nucleosomes and histones and for performing its main physiological functions, although deleting distinct parts of the acidic domain produced different, partially additive phenotypes, indicating specific but diffusely distributed functions (Figs 2C, 3). We therefore propose that the acidic domain is important for substrate recognition, and that the N-terminal region of this domain may have a dispersed, non-specific electrostatic role while the C-terminal portion contributes more to interaction with other factors like Spt6p.

The interaction with Spt6p is not required for Tom1p activity, as indicated by our finding that deletion of the binding site did not cause the growth defects associated with full loss of Tom1p. Interaction with Spt6p is therefore more likely required to optimize some function of Tom1p, possibly by regulating when or where it should be active on a subset of targets. The interaction of Spt6p^tSH2^ with Rpb1p has a similarly non-essential but advantageous function (Sdano et al. 2017, Connell et al. 2022). In that case, Spt6p^tSH2^ appears to collaborate with the Paf1 complex to make a stable contact with the Rpb1p linker (Ellison et al. 2023), suggesting that the tSH2 binding sites are typically occupied by Rpb1p during elongation. The pattern of Tom1p occupancy (low in promoters, high in gene bodies) suggests it might be tethered to elongation complexes, while the effect of loss of Tom1p on nucleosome occupancy (decreased in promoters, increased in gene bodies) suggests that it might be stabilizing nucleosomes in promoters and destabilizing them in gene bodies. Taken together, these observations suggest that Tom1p associates with Spt6p in promoters before the elongation complex forms and that this spares free histones from degradation, while a looser or different association of Tom1p with the elongation complex enhances its occupancy in gene bodies where it is more prone to promoting histone turnover. Other models are possible and will require further testing.

The 259 residues of the acidic domain of Tom1p lack density in our structural analysis, presumably due to being disordered. One possibility is that this ∼29 kDa domain occupies the solenoidal basket, perhaps extending above the basket rim, and that it retains basic substrate proteins in the central region of the solenoid. It is also possible that the shift from the closed ring to the open spiral form is important for function. Mutations expected to destabilize closure of the ring conformation were reported to compromise E3 ligase activity of human HUWE1 (Hunkeler et al. 2021) and similar mutations of *Nematocida* HUWE1 were found to destabilize the protein (Grabarczyk et al. 2021). It has also been proposed that opening of the ring releases the HECT domain into a more flexible environment that allows its active site to turn towards substrates bound to N-terminal regions of the protein (Hunkeler et al. 2021) (Grabarczyk et al. 2021). The functional relevance of the ring vs spiral conformations, and the possibility of functional transition between the two states, is an important topic for future studies.

In summary, this work reveals an unexpected role for Tom1p/HUWE1 family members in maintaining the appropriate pools of partially assembled nucleosomal components in different contexts to maintain differential chromatin architecture in promoters and transcribed regions. One possibility is that the interaction with Spt6p serves to localize Tom1p/HUWE1 to these regions. It is also possible that the Spt6p^tSH2^-Tom1p interaction competes with or modulates the Spt6p^tSH2^-Rpb1p interaction, thereby providing information to this hub to assist in coordinating progression of the elongation complex. The structures provided here will be a basis of future work to investigate these ideas.

## METHODS

### Strains/constructs

Yeast strains (Table S1) were isogenic with the A364a background and were constructed using standard methods. The *tom1-Δ*0 allele was transferred from the standard deletion collection (Research Genetics; Brachmann et al. 1998) into the A364a background using primers TF1654, TF1655 (Table S2) to construct strain 9731 (Table S1), which was used to construct additional strains in this background using standard genetic methods.

The pCORE cassette (Storici et al. 2001) was integrated into the *TOM1* locus in the promoter region (primers TF1658, TF1659), after residue K1074 (primers TF2111, TF2112), and after S1943 (primers TF1714, TF1715). These strains were then used to swap the native *TOM1* promoter with the *GAL1* promoter, to integrate tags, and to create various internal deletions of the acidic domain using the primers listed in Table S2.

The promoter region of *TOM1* was replaced with the *GAL1* promoter using the delitto perfetto method (Storici et al. 2001), then a PreScission site-Protein A sequence was fused in-frame to the C-terminus of the Tom1p ORF (strain 9750, Table S1). A 20-residue encoding SGGGGGGSRHHHHHHHHHHHH was fused in-frame between K1074 and D1075 in a loop region identified in the Tom1p cryo-EM structure, also using the delitto perfetto method (Storici et al. 2001). Primers for these constructions are listed in Table S2.

### Plasmids

#### Strep-FLAG-Spt6p^tSH2^ construct

The coding sequence for Spt6p(1223-1451) was amplified from genomic DNA using PCR primers TF0950 and TF0951 (Table S2). The product was digested with Nde1 and BamH1, then ligated with pTF198 (YEp_LEU2_Gal1) digested with the same enzymes to create pTF254 with a FLAG epitope fused to the N-terminus of the Spt6p tSH2 domain. *Kpn1* and *Xho1* sites were introduced between the FLAG epitope and the Spt6p^tSH2^ coding sequence using site-directed mutagenesis with primers MS1+/-to create pTF254-KX. A twin-Strep-tag sequence (IBA) flanked by Kpn1 and Xho1 sites was then constructed by annealing primers MS351, MS352 and this was inserted into pTF254-KX to yield pMS14 (CPH3375) expressing an N-terminal streptavidin-FLAG tandem affinity tag (SF-TAP; Gloeckner et al. 2007) fused to Spt6p^tSH2^, followed by an in-frame C-terminal SV40 nuclear localization signal, under control of the *GAL1* promoter.

#### GST-Spt6p^tSH2^ construct (WT and -R, -KK, -R,KK)

Plasmids expressing GST-tagged Spt6p were produced by inserting the Spt6p(1223-1451) coding sequence into the pDEST15 vector (Thermo Fisher Scientific). Mutations in the tSH2 domain were introduced using site-directed mutagenesis (Sdano et al. 2017).

### Protein expression and purification

All protein purification steps were performed at 4°C, unless otherwise stated.

#### Strep-FLAG-Spt6p^tSH2^

Yeast transformed with plasmid pMS14 [SF-Spt6p^(1223-1451)^ or pTF198 (empty) were cultured in two liters of synthetic media containing glycerol and lactate in baffled flasks at 30 °C. When the OD^600^ reached ∼1, expression was induced by the addition of galactose to 0.55% and incubation was continued overnight. Cells were harvested by centrifugation and lysed under liquid nitrogen using a freezer mill. Subsequent steps were performed at 4°C. Yeast powder was suspended in an equal volume of lysis buffer (100 mM Tris•Cl pH 7.5, 300 mM NaCl, 10% glycerol, 1 mM DTT, 0.1% Tween-20, 1x protease inhibitors [0.5 μg/mL aprotinin, 0.5 μg/mL leupeptin, 0.7 μg/mL pepstatin, 167 μg/mL PMSF], 1x phosphatase inhibitors [2 mM NaVO_4_, 2 mM BGP, 2 mM NaF], 6 μg/mL DNAse I, 2 μg/mL RNAse A) and clarified by centrifugation at 107,000 RCF for 30 minutes. 120 μg avidin was added to the clarified lysate and incubated for 15 minutes to block biotinylated proteins. Lysates were clarified further by ultracentrifugation at 107,000 RCF for 1 hour. The supernatant was applied to 100 μl Strep-Tactin® Superflow resin (IBA) and incubated for 1 hour. The resin was washed with 10 column volumes of wash buffer (50 mM Tris•Cl pH 7.5, 150 mM NaCl, 5% glycerol, 0.1% Tween-20, 1x protease and phosphatase inhibitors, 6 μg/mL DNAse, 2 μg/mL RNAse A) and eluted with buffer containing 2 mM desthiobiotin. The eluate was applied to 50 μl anti-FLAG^®^ M2 affinity gel (Sigma-Aldrich) and incubated for 1 hour. The resin was washed 3 times with 100 μl wash buffer and eluted with 150 μl wash buffer containing 200 μg/mL FLAG peptide. Proteins were separated by SDS-PAGE on NuPAGE™ Novex™ 4-12% Bis-Tris Protein Gels (Invitrogen) in MOPS running buffer, and proteins were visualized by staining with Coomassie Blue dye.

#### GST-Spt6p^tSH2^

GST-Spt6p^tSH2^ proteins were expressed and purified as described previously (Close et al. 2011), except the lysate was applied to glutathione agarose resin (Pierce) and the proteins were not cleaved but instead were eluted with 10 mM glutathione to leave the GST tag intact.

#### Tom1p and Tom1p^ΔAD^

*S. cerevisiae* strains 9750 (intact Tom1p-PrtA) or 10232 (Tom1p^ΔAD^-PrtA) was grown on an orbital shaker in non-baffled flasks in YP media containing glycerol and sodium lactate to saturation at 30 °C, and protein expression was induced overnight via addition of galactose to 0.5% w/v. Cells were harvested by centrifugation at 5,000 RCF, lysed under liquid nitrogen using a freezer mill, and stored at -80 °C. 40 g pulverized yeast lysate were added to 150 mL total lysis buffer (containing 50 mM Tris•Cl pH 7.5, 500 mM NaCl, 10% glycerol, 0.1% Tween-20, 1 mM DTT). 167 ug/mL PMSF, 0.7 ug/mL aprotinin, 0.7 ug/mL leupeptin, and 0.9 ug/mL pepstatin as protease inhibitors, plus 2 mM BGP and 2 mM NaF as phosphatase inhibitors were added to the initial lysate solution. Cell lysate was stirred vigorously at 4 °C for 10 minutes, vortexed briefly for 1 second, and clarified via centrifugation at 18,000 RCFfor 35 minutes.

Clarified lysate was added to 2.3 mL of a 50% IgG Sepharose 6 Fast Flow resin (GE Healthcare) slurry and incubated on a roller for 1 hour. Resin was washed 4x with 10 mL of suspension buffer and with 4x 10 mL of a wash buffer containing 50mM Tris•Cl pH 7.5, 150mM NaCl, 10% glycerol, 0.1% Tween-20, 1mM DTT. Resin was then incubated for 2 hours with 400 uL of 1mg/mL PreScission protease (PP) in 15 mL of wash buffer additionally containing 1% Tween-20, 0.5 mM EDTA•Na2, plus protease and phosphatase inhibitors as above. Eluent was combined with an additional 2x 20 mL of wash buffer and subsequently applied to a 5 mL HiTrap Q (GE Healthcare) ion-exchange column equilibrated in a buffer containing 25 mM Tris•Cl, 5% glycerol, 0.5mM EDTA•Na2, and 2mM 2-mercaptoethanol, and eluted over a gradient from 0 mM to 1 M NaCl. Tom1p eluted at approximately 500 mM NaCl. Fractions containing Tom1p were then pooled, concentrated, and loaded onto a Superdex S6 size exclusion column (GE Healthcare) equilibrated in 5% glycerol, 50 mM Tris•Cl pH 7.5, 150 mM NaCl, and 2 mM DTT. Fractions containing Tom1p were pooled and concentrated.

#### Tom1p^His^

*S. cerevisiae* strain 10293 (Tom1p^His^) was purified as above, with the following differences: DTT was replaced by 2.5 mM 2-mercaptoethanol in the lysis buffer, IgG beads were replaced by 3.5 mL Nickel beads. Wash buffer contained 25 mM Tris•Cl pH 6.8, 125 mM NaCl, 5 mM imidazole, and no DTT. Elution buffer contained 25mM Tris•Cl pH 6.8, 125 mM NaCl, 10% glycerol, 0.1% Tween-20, 500 mM imidazole, 0.5 mM EDTA•Na2, 10 mM 2-mercaptoethanol, plus protease and phosphatase inhibitors as above. Resin was washed 3x with 2 CV lysis buffer, 1x 2 CV lysis buffer containing 0.2ul RNAse A first incubated for 5 minutes, 3x 2 CV wash buffer, and 4x 2 CV Elution buffer. Before application to HiTrap Q column, eluent was dialyzed overnight at 4 °C into 2L of an elution buffer containing 7.5% glycerol, no imidazole, and 2.5 mM 2-mercaptoethanol.

### Far-Western blots

Far-western blots were performed essentially as described previously (Wu et al. 2007). Either 20 μL of the FLAG elution described above or 1.1 pmol purified Tom1p were loaded to a NuPAGE™ Novex™ 4-12% Bis-Tris Protein Gel (Invitrogen) and electrophoresed in MOPS buffer. Proteins were transferred to nitrocellulose membranes for 70 minutes at 100 V. After a 1-hour in blocking buffer (1xTBS, 3% milk powder) the membrane was incubated in GST-Spt6p^tSH2^ protein in binding buffer (100 mM NaCl, 20 mM Tris•Cl, 10% glycerol, 3% milk powder, 0.2% EDTA•Na2, 0.1% Tween-20, 0.1% DTT, 50 nM GST-Spt6p^tSH2^) overnight at 4°C. Anti-GST primary antibody was used at 1:1,000 (GE Healthcare 27-4577-01) and secondary antibody was conjugate to Alexa 680 nm dye at 1:10,000, washing with 1xTBS between steps.

For phosphatase treatment, purified Tom1p was diluted to 300 nM in phosphatase buffer (50 mM Tris•Cl pH 7.5, 100 mM NaCl, 10 mM MgCl_2_, 5% glycerol, 0.1% Tween-20, 1 mM DTT, 1x protease inhibitors) and incubated with 0.3 U/μl CIP (New England BioLabs) for 1 hour at room temperature prior to loading onto a gel.

### Limited proteolysis

100 μg of purified Tom1p was incubated with 100 ng trypsin in 200 μL sizing buffer at room temperature. Time points were collected at 30 seconds, and 1, 3, 10, 20 and 30 minutes. Reactions were quenched by the addition of aprotinin, leupeptin, PMSF, and SDS-loading dye followed by boiling for 3 minutes. Time points were resolved by SDS-PAGE on NuPAGE™ Novex™ 4-12% Bis-Tris Protein Gels (Invitrogen) in MOPS running buffer, and proteins were visualized by Coomassie staining or transferred to a membrane for far western blotting.

### Mass spectrometry

LC/MS/MS analyses of the Spt6p Tom1p-binding domain visualized from limited proteolysis and identification of the Tom1p protein (Figure 2B) were performed as described (Sdano et al. 2017). Proteomics mass spectrometry analysis was performed at the Mass Spectrometry and Proteomics Core Facility at the University of Utah.

### Western blots

Western blots were performed as previously described except 60 µg of total protein was loaded per lane (McCullough et al. 2015), and antiserum obtained from a rabbit injected with purified Tom1p protein was used (Covance). Specificity of the antiserum was validated using strains lacking Tom1p (see Fig. 2C).

### Cryo-EM specimen grid preparation

Specimens were prepared on Quantifoil R 2/2 Cu300 grids after glow discharge for 1 minute at 25 mA. For Tom1p^ΔAD^ samples, Tom1p^ΔAD^ was concentrated to 8.1 mg/mL in 0.01% CHAPS and 0.05% NP-40. 2.5 μL of sample was applied to grids and blotted for 2.5 seconds before vitrification by plunging into liquid ethane. For Tom1p^ΔAD^ samples without detergent, protein was concentrated to 0.44 mg/mL and blotted for 1.5 seconds before vitrification. For Tom1p^His^, protein samples were prepared as for Tom1p^ΔAD^ without detergent and concentrated to 0.5 mg/mL. 2.5 μL of sample was added to grids and blotted for 2.5 seconds before vitrification.

### Cryo-EM data collection

A total of 2,865 movies were recorded from Tom1p^ΔAD^ grids without detergent at 81,000x magnification on a 300 kV Titan Krios G3 electron microscope with a K3 direct detector (nominal resolution after 2x binning is 1.06Å/px). Electron exposure was 47 (e-/Å^2^) over a total of 40 frames. A total of 4,371 movies were recorded from Tom1p^ΔAD^ with detergent grids using the same electron microscope and collection parameters. A total of 3,630 movies were recorded from Tom1p^His^ grids using the same electron microscope and collection parameters, with a total electron exposure of 40 (e-/Å2) over 40 frames.

### Cryo-EM data processing

Cryo-EM data processing was performed in CryoSPARC (Punjani et al. 2017). Movies were motion-corrected for all 3 datasets using CryoSPARC Patch Motion Correction. CTF estimation was performed on motion-corrected micrographs using CryoSPARC Patch CTF with default parameters applied.

#### Dataset #1: Tom1p^ΔAD^-no detergent

Initial ‘blob’ auto-picking, 2D classification, *ab initio* reconstructions and heterogeneous refinements led to 3 refined volumes. Templates were low-pass filtered to 17 Å creating 25 equally spaced templates each for a total of 75 templates. These templates were used to template-pick from 2,139 micrographs selected from the original 2,865 by CTF resolution (max 5.6 Å CTF estimated resolution). An initial set of 792,807 particles was filtered by 2D classification to 513,104 particles, followed by *ab initio* volume creation to generate 5 classes. Three of the classes were removed as junk and the remaining particles subjected to additional 2D and 3D classification to yield 235,202 particles that were combined with particles from the Tom1p^ΔAD^-detergent dataset for downstream processing.

#### Dataset #2: Tom1p^ΔAD^-detergent

4,356 of the 4,371 micrographs from the Tom1p^ΔAD^-detergent dataset were selected after inspection of CTF estimation, and the 75 templates from Dataset #1 (above) were used to select 544,686 template-picked particles. Multiple rounds of 2D and 3D classification were used to remove junk and any open- or intermediate-conformation particles, which narrowed the selection to 296,347 particles.

#### Combined Datasets #1/#2: Tom1p^ΔAD^

The separately processed dataset #1 and #2 particles were combined for a total of 531,549 starting particles for further processing and refinement. A round of 2D classification resulted in selection of 477,889 particles, which were Fourier cropped from a box size of 680 to 320 pixels for further processing. Two-component 3DVA was performed after map refinement of the combined dataset with a filter resolution of 4 Å. 3DVA display was subsequently run to give 8 intermediates along the first principal component at a filter resolution of 5 Å. The top 4 intermediates with the highest number of particles were combined to yield 462,711 and refined using non-uniform refinement to an estimated overall resolution of 3.07 Å.

#### Dataset #3: Tom1p^His^

3,522 of 3,630 micrographs were selected for particle picking from grids containing Tom1p^His^. Initial blob picks were curated via 2D-classification and 3D ab initio jobs to create 60 2D templates for template-picking. 1,434,189 template-picked particles (lowpass filter 15 Å) were narrowed through 6 rounds of 2D classification to give 201,930 particles, which were input to *ab initio* volume creation with 4 classes. As outlined below, 41,253 particles from one of these classes were used to refine the open/intermediate conformations and 160,667 total particles from the remaining 3 classes were combined for further refinement of the closed conformation map.

Closed conformation: The selected 160,667 particles were combined for 4-class 3D *ab initio* volume creation to remove one junk class. 2D classification on the remaining particles was used to remove 6,726 additional junk/low-resolution particles, and ab initio and NU-refinement was performed on the final 142,020 particles, resulting in a closed-conformation map at an estimated overall resolution of 3.97 Å.

Open/intermediate conformations: The selected 41,253 particles were assessed for *ab initio* reconstruction in 4 classes. NU-refinement was performed on 2 of these classes to give an intermediate-conformation map at 7.79 Å resolution from 12,397 particles and an open-conformation map at 7.10 Å from 16,098 particles.

### Model building and refinement

Initial model building into the Tom1p^ΔAD^ closed-conformation map was done in Coot v.0.9.6 via Xquartz 2.8.2 as part of the CCP4-7.1 Program Suite (Emsley et al. 2010). In parallel with *ab initio* model building, portions of the sequence were predicted iteratively for the entire protein, approximately 200-500 residues at a time, using the AlphaFold2 algorithm employed by ColabFold v.1.0 and accessed via a Google Colab Jupyter Notebook (Mirdita et al. 2022). The top models predicted for each segment of sequence were compared with the corresponding portions of the manually built model and used for validation of residue placement and additional model refinement. The HECT domain model was built by fitting the x-ray structure of the Tom1p HECT domain to the density. The model was then subject to Real Space Refinement via Phenix v.1.19.1.(Afonine et al. 2018) using secondary structure restraints and a target bond/angle RMSD of 0.1 Å and 1.56°, respectively. Statistics are summarized in Table S3. All model and map figure and movie generation were done via Chimera (Pettersen et al. 2004) or ChimeraX (Meng et al. 2023).

### Structural Cα alignment

The aligned Cα RMSD between the full sequence of yeast Tom1p and human HUWE1 structures was calculated using the Matchmaker function integrated in ChimeraX, applying the Needleman-Wunsch alignment algorithm, BLOSUM-62 similarity matrix (Goddard et al. 2018).

### **X-** ray structure of Tom1p HECT domain

An His-MBP-TEV-Tom1(HECT: 2850-3268) was expressed and purified according to standard protocols. Briefly TEV protease cleavage occurred overnight in dialysis to 15 mM Tris pH 7.5, 100 mM NaCl, 5% glycerol, 0.5 mM EDTA pH 8.0, 2 mM BME and then run on an S200 size exclusion column, from which it eluted as a monomeric peak. The protein was concentrated to 15 mg/mL prior to crystallization. Data were collected at SSRL beamline 12-2 with wavelength 0.97946 Å. Images were processed with XDS, Aimless, and Truncate in CCP4 (Agirre et al. 2023). (Table S4). The model was built and refined in Coot (Emsley et al. 2010) and Phenix (Afonine et al. 2018) to 3.3Å. The modeled structure includes residues 2852-3261, except residues 2904-2908, 2991-2996, 3079-3090 and 3064-3065. The structure was refined at 3.3Å resolution. The structure closely resembles the HUWE1 HECT domain structure (PDBcode:5lp8) with an overall RMSD of 0.9Å and only two areas of divergence. The HUWE1 structure contains a two ordered β-strands in the middle domain residues 4172-4196 whereas the Tom1p HECT contains disordered loop residues between 3078-3091.

Maps and models have been deposited to the EMDB and PDB respectively. Tom1^ΔAD^ cryoEM maps closed (EMDB:XXXX/PDB:XXXX), intermediate map (EMDB:XXXX), open map (EMDB:XXXX), and the x-ray structure of the Tom1 HECT (PDB:XXXX). Plasmids were deposited to Addgene.org.

## ACKNOWLEDGMENTS

We gratefully acknowledge the support and resources of the University of Utah Center for High Performance Computing. We thank the University of Utah HCI Cores for ChIP, MNase, and nucleosome profiling, and the University of Utah HSC cores for DNA sequencing, Mass Spectrometry and Proteomics, and Electron Microscopy. We thank Dr. David Belnap for assistance with EM data collection and Dr. Frank Whitby for x-ray data collection. Use of the Stanford Synchrotron Radiation Lightsource, SLAC National Accelerator Laboratory, is supported by the U.S. Department of Energy, Office of Science, Office of Basic Energy Sciences under Contract No. DE-AC02-76SF00515. The SSRL Structural Molecular Biology Program is supported by the DOE Office of Biological and Environmental Research, and by the National Institutes of Health, National Institute of General Medical Sciences (including P41GM103393). Core Facility Mass spectrometry equipment was obtained through NIH Grant S10 OD018210. CryoEM instrumentation was obtained with support from the Arnold and Mabel Beckman Foundation. This work was supported by NIH grants U54AI170856 (CPH) and GM064649 (TF). The contents of this publication are solely the responsibility of the authors and do not necessarily represent the official views of NIH.

