## Supplemental Information 1 for "Tom1p ubiquitin ligase structure, interaction with Spt6p, and function in maintaining normal transcript levels and the stability of chromatin in promoters"

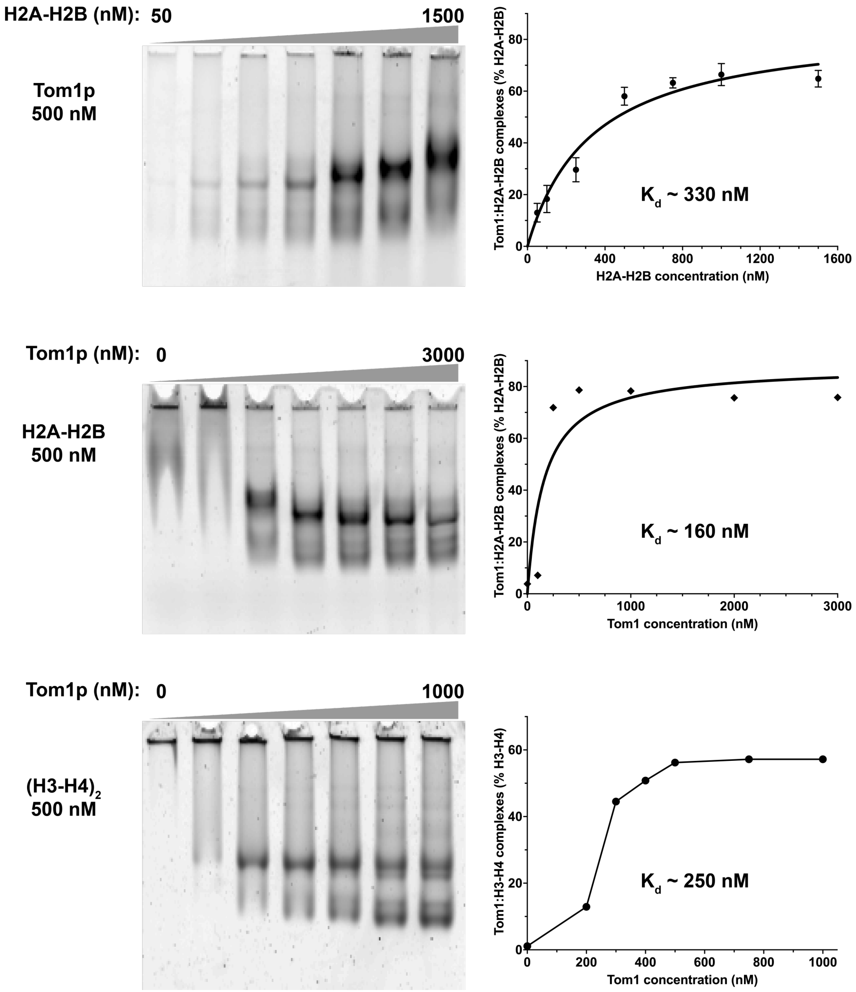

**Figure S1. Affinity of Tom1p for histones determined by EMSA**

Top: H2A-H2B dimers were titrated (50-1500 nM) with Tom1p (500 nM) and complexes were detected as in Figure 4 as the fraction of H2B signal in the prominent band; also see (McCullough et al. 2018). Samples were tested in quadruplicate with one set shown, and the average (+/- SD) used to calculate the binding affinity in Prism 9.

Middle: Tom1p was titrated (0-3000 nM) with H2A-H2B (500 nM) and complexes were detected by EMSA as above. Quantitation of this and similar experiments typically produced Kd values from 100-200 nM (160 nM for the experiment shown).

Bottom: As in the middle panel, except using 500 nM $(H3\text{-}H4)_2$ tetramers, which are recovered inefficiently in this assay unless bound to Tom1p.

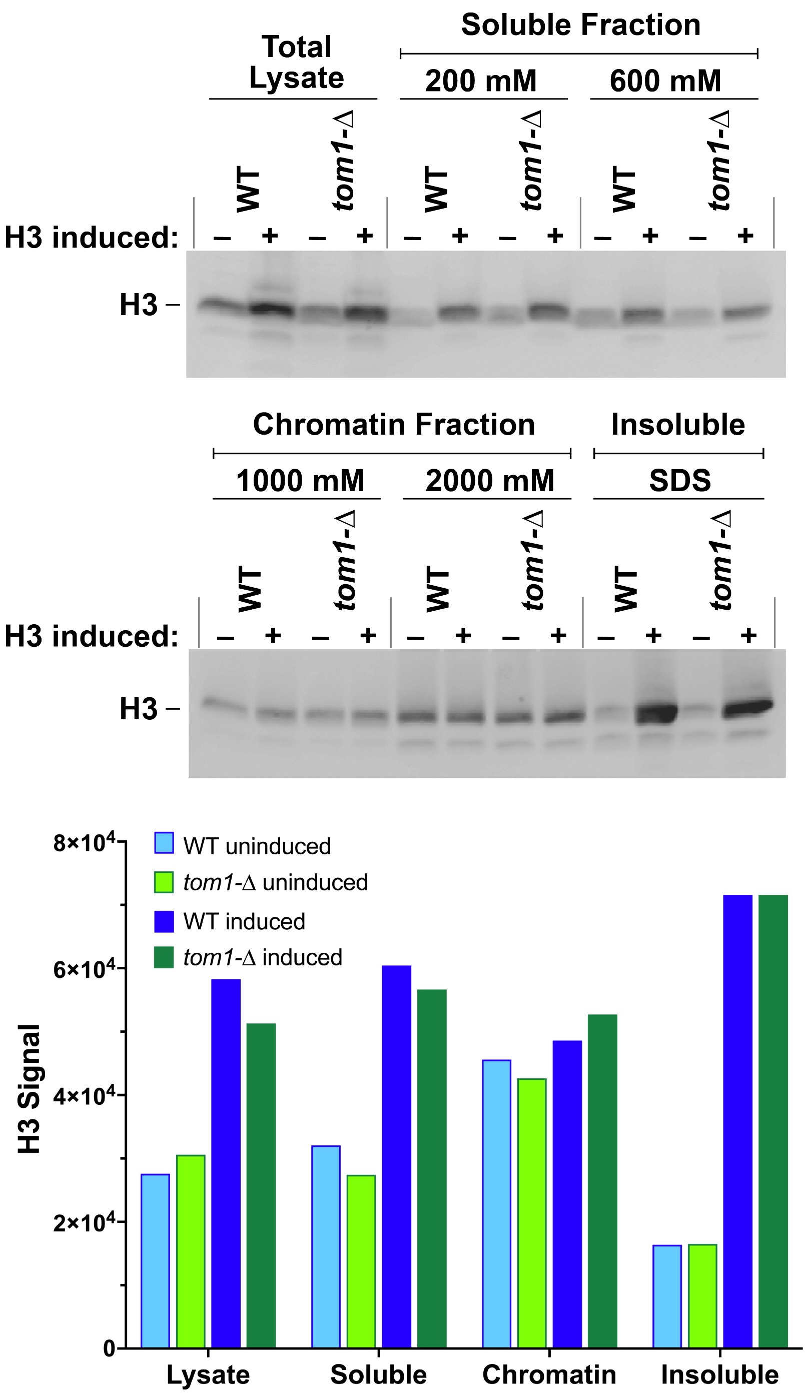

**Figure S2. Loss of Tom1p does not significantly affect the level of histone H3 in soluble or chromatin-associated pools.**

Tom1p has been implicated in global turnover of histones in yeast (Singh et al. 2009) but we found that cells lacking Tom1p had normal levels of the histone H3 (panel A, total lysate samples). To probe this further, we overexpressed H3 by placing the *HHT2* ORF under control of the *GAL1* promoter (we found that overexpression at the level of protein required removal of the 3' untranslated region of *HHT2* and substitution of this region with the terminator from

*ADH1*; not shown). We then asked whether the turnover of excess H3 protein required Tom1p, either from a soluble pool (presumably, H3 ready for assembly of chromatin) or from the chromatin pool itself. Instead, we found that the bulk of the excess H3 produced by overexpression of *HHT2* was found in an insoluble fraction that was only recovered by denaturation with SDS sample buffer ("insoluble" fraction). While overexpression also increased the level of soluble H3, in no case did we see significantly higher levels of H3 in cells lacking Tom1p, suggesting that Tom1p was not required for degrading excess H3 whether expression was normal or elevated. Similar results were obtained with two technical replicates of the samples shown, and from several biological replicates using variations of the extraction protocol (not shown).

H3 levels were measured as follows: Strains 8127-7-4 (WT) and 9731-1-3 (*tom1*-Δ) were transformed with plasmid pZC23 (YEp *URA3 GAL1p HHT2* ORF *ADH1* terminator) and grown to an OD of about 0.4 in synthetic medium lacking uracil with 2% raffinose as the carbon source at 30° C. About 3 X $10^7$ cells were harvested as the uninduced samples and held on ice. Galactose was added to the remaining culture to a final concentration of 1% and growth was continued for 3 hours to a final OD of about 1.0. About 3 X $10^7$ cells were harvested as the induced samples. Cells were collected by centrifugation, washed twice with 25 mL of 100 mM Tris•HCl (pH 9.4) at 0 °C, then suspended in 10 mL of the same buffer with 10 mM dithiothreitol and incubated at 0 °C for 15 minutes. Cells were collected by centrifugation, washed once in spheroplasting buffer (20 mM HEPES pH 7.4, 1.2 mM Sorbitol), then suspended in 2 mL of the same buffer with 50 µg/mL Zymolyase 100T (Fischer) and incubated at 30 °C for 10 minutes (>90% of a test aliquot lysed when added to water). Cells were collected by centrifugation at 4 °C, washed twice with 2 mL wash buffer (20 mM Tris•HCl pH 7.4, 20 mM KCl, 1 M sorbitol, 0.1 mM spermine, 0.25 mM spermidine), then suspended in 1 mL of lysis buffer (20 mM Tris•HCl pH 7.4, 200 mM NaCl, 1 mM spermine, 2.5 mM spermidine, 1% Triton X-100). After incubation at 0 °C for 18 minutes, 200 µl of the suspension (~6 X $10^6$ cell equivalents) was removed and mixed with 50 µl of 5X SDS sample buffer as the total lysate fraction. 700 µl of the remaining lysate (~2.1 X $10^7$ cell equivalents) was centrifuged at 15,000 X g for 15 minutes at 4 °C, the supernatant was removed and mixed with 5X SDS sample buffer as the 200 mM NaCl fraction. The pellet was suspended in extraction buffer (Tris•HCl pH 7.4, 600 mM NaCl), incubated for 5 minutes at 22 °C, then centrifuged as above. The supernatant was removed as the 600 mM fraction, prepared for SDS PAGE as above, then this procedure was repeated with increasing NaCl concentrations to generate 1M and 2M extracts. Finally, the remaining pellet was solubilized in 750 µl of 1X SDS sample buffer as the insoluble fraction. Aliquots were analyzed by SDS PAGE with samples from the total lysate (representing about 3 X $10^5$ cell equivalents), and extractions (about 4.5 X $10^5$ cell equivalents). Western blot analysis was performed using antibodies generated against a peptide from yeast histone H3 injected into a rabbit (Covance) followed by secondary detection with a goat anti-rabbit IgG attached to Alexa Fluor plus 800 (Li-Cor). After quantitation (Odyssey imager, Li-Cor), signals from the 200 mM and 600 mM extracts were summed as the soluble H3, and signals from the 1M and 2M NaCl extracts were summed as the chromatin fraction.

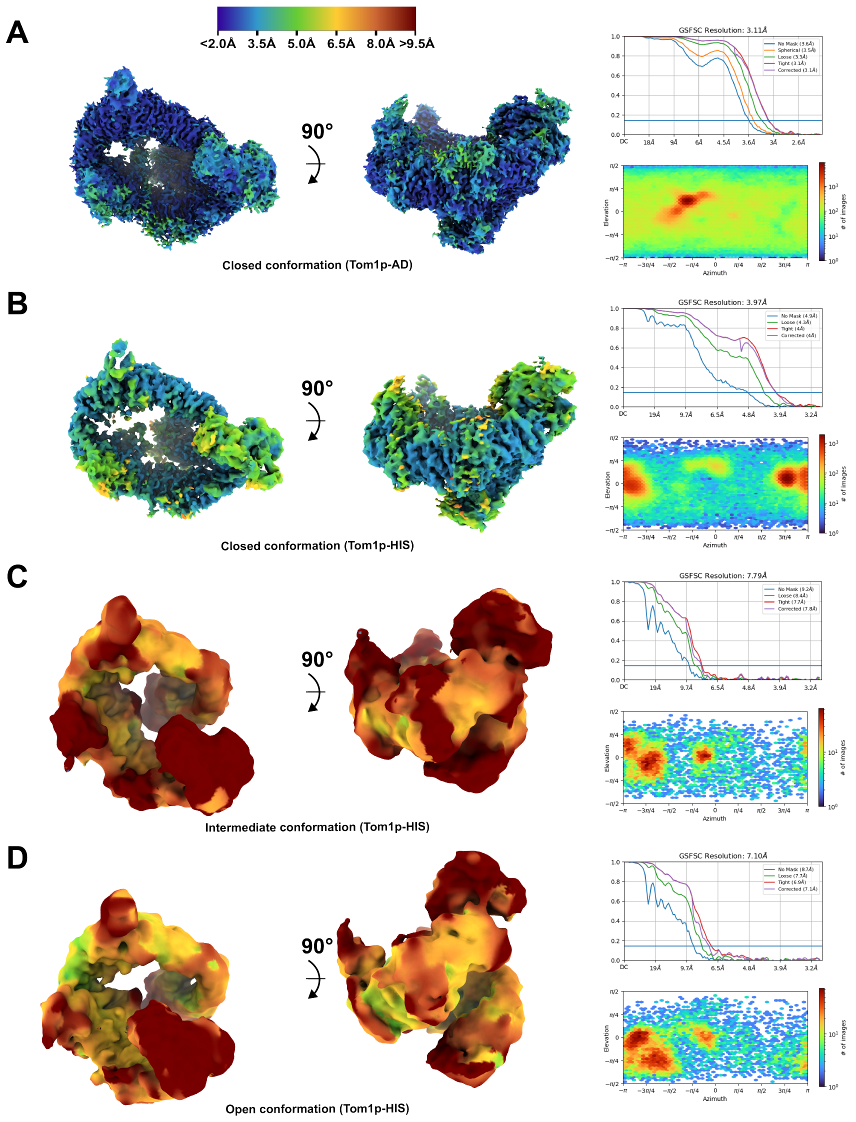

**Figure S3. Tom1p resolution and particle orientation distribution.**

**A)** Tom1p$^{\Delta AD}$ closed conformation. **B)** Tom1p$^{His}$ closed. **C)** Tom1p$^{His}$ intermediate. **D)** Tom1p$^{His}$ open. Two orientations of maps (same as Fig. 6) show local resolution, aligned from residues 1-1315. Gold-standard FSC resolution estimation and plots of orientation distribution are also shown.

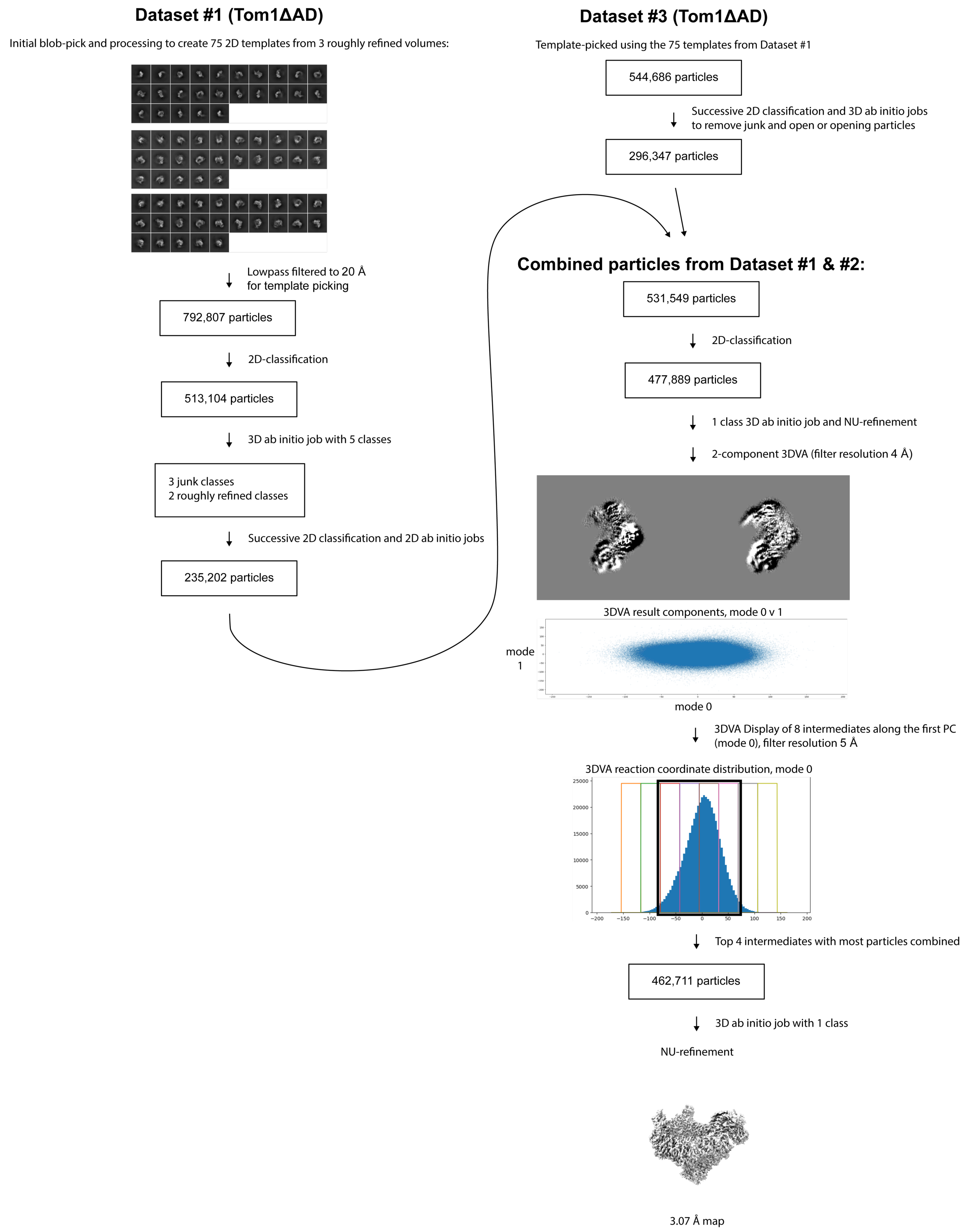

**Figure S4 Cryo-EM processing workflow for the Tom1p$^{\Delta AD}$ structure**

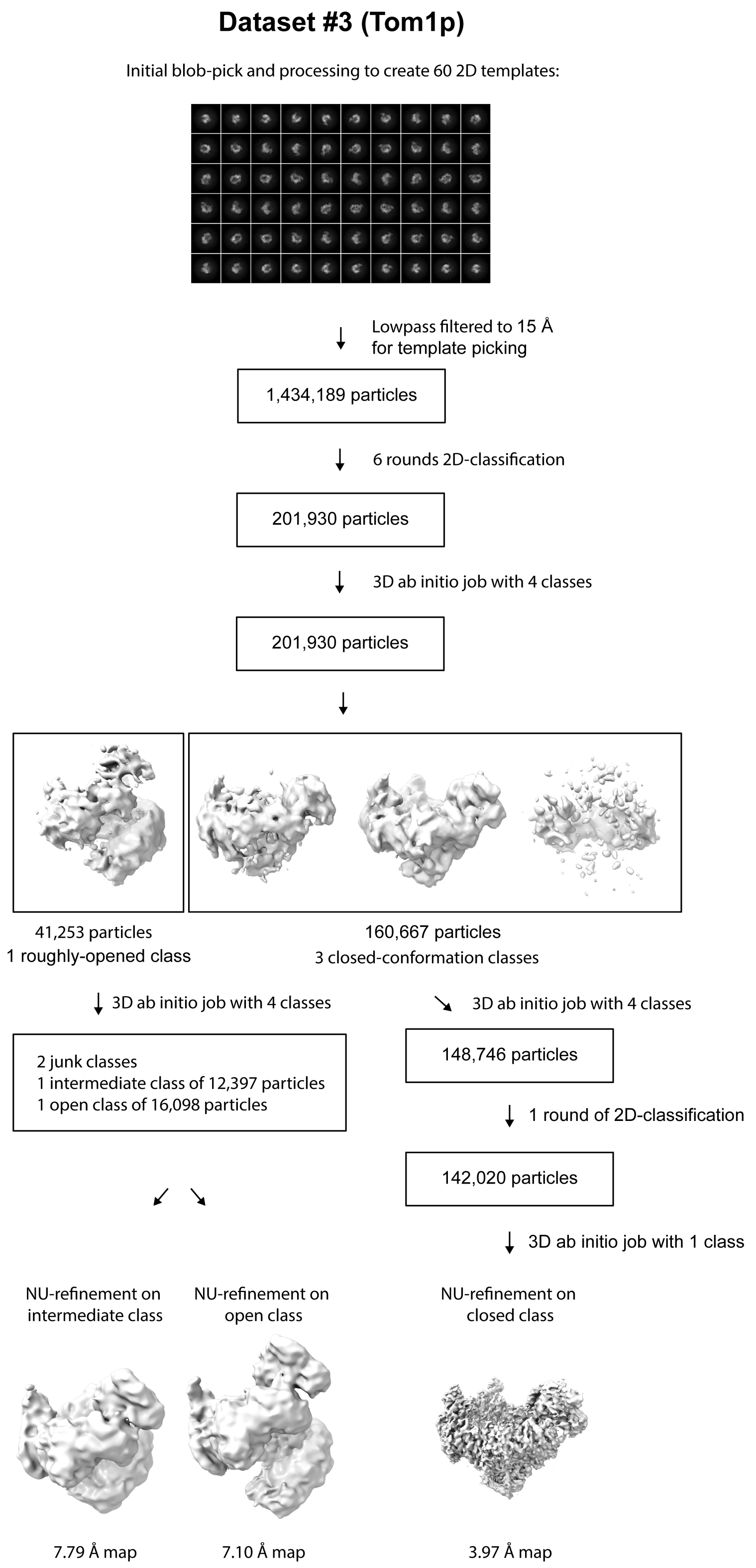

**Figure S5. Cryo-EM processing workflow for the Tom1p$^{His}$ structures**

**Table S1. Yeast strains**

| | | |
|---|---|---|
| Strains used in Figure 2 | | |
| Name | Label | Full Genotype |
| 10228-2D | *TOM1* | *MATa ura3-Δ0 leu2-Δ0 trp1-Δ2 his3 lys2-128∂ rpb1-LMFSPLV1490KMKRRRK(+100, HphMX)* |
| 10243-3D | *tom1-Δ0* | *MATa ura3-Δ0 leu2-Δ0 his3 lys2-128∂ rpb1-LMFSPLV1490KMKRRRK(+100, HphMX) tom1-Δ0(::KanMX)* |
| 10236-7D | *tom1-intΔ2 (Δ1873-2131)* | *MATa ura3-Δ0 leu2-Δ0 trp1-Δ2 his3 lys2-128∂ rpb1-LMFSPLV1490KMKRRRK(+100, HphMX) tom1-Δ2(1873-2131)(+50, KanMX)* |
| 10238-11C | *tom1-intΔ4 (Δ1873-2022)* | *MATa ura3-Δ0 leu2-Δ0 his3 lys2-128∂ rpb1-LMFSPLV1490KMKRRRK(+100, HphMX) tom1-Δ4(1873-2022)(+50, KanMX)* |
| 10239-3C | *tom1-intΔ5 (Δ1873-1972)* | *ura3-Δ0 leu2-Δ0 his3 lys2-128∂ rpb1-LMFSPLV1490KMKRRRK(+100, HphMX) tom1-Δ5(1873-1972)(+50 KanMX)* |
| 10270-10A | *tom1-intΔ9 (Δ1973-2131)* | *MATa ura3-Δ0 leu2-Δ0 his3 lys2-128∂ rpb1-LMFSPLV1490KMKRRRK(+100, HphMX) tom1-Δ9(1973-2131)(+50 KanMX)* |
| 10291-1D | *tom1-intΔ10 (Δ1998-2059)* | *MATa ura3-Δ0 leu2-Δ0 trp1-Δ2 his3 lys2-128∂ rpb1-LMFSPLV1490KMKRRRK(+100, HphMX) tom1-Δ10 (1998-2059)(+50 KanMX)* |
| 10292-3D | *tom1-intΔ11 (Δ2031-2063)* | *MATa ura3-Δ0 leu2-Δ0 trp1-Δ2 his3 lys2-128∂ rpb1-LMFSPLV1490KMKRRRK(+100, HphMX) tom1-Δ11 (2031-2063)(+50 KanMX)* |
| Strains used in Figure 2S | | |
| 10228-2D | *TOM1* | *MATa ura3-Δ0 leu2-Δ0 trp1-Δ2 his3 lys2-128∂ rpb1-LMFSPLV1490KMKRRRK(+100, HphMX)* |
| 10243-3D | *tom1-Δ0* | *MATa ura3-Δ0 leu2-Δ0 his3 lys2-128∂ rpb1-LMFSPLV1490KMKRRRK(+100, HphMX) tom1-Δ0(::KanMX)* |
| 10291-1D | *tom1-intΔ10 (Δ1998-2059)* | *MATa ura3-Δ0 leu2-Δ0 trp1-Δ2 his3 lys2-128∂ rpb1-LMFSPLV1490KMKRRRK(+100, HphMX) tom1-Δ10 (1998-2059)(+50 KanMX)* |
| 10292-3D | *tom1-intΔ11 (Δ2031-2063)* | *MATa ura3-Δ0 leu2-Δ0 trp1-Δ2 his3 lys2-128∂ rpb1-LMFSPLV1490KMKRRRK(+100, HphMX) tom1-Δ11 (2031-2063)(+50 KanMX)* |
| Strains used in Fig 3 | | |
| 10144-6C | *TOM1* | *MATa ura3-Δ0 leu2-Δ0 trp1-Δ2 his3 lys2-128∂ TOM1(+50, KanMX)* |

| | | |
|---|---|---|
| 9731-1-3 | *tom1-Δ0* | *MATa ura3-Δ0 leu2-Δ0 trp1-Δ2 his3 lys2-128∂ tom1-Δ0(::KanMX)* |
| 9825 | *tom1-S1943-pCORE-D1944* | *MATa ura3-Δ0 leu2-Δ0 trp1-Δ2 his3 lys2-128∂ tom1-S1943(::URA3 Kl, KanMX)* |
| 10236-1A | *tom1-intΔ2 (Δ1873-2131)* | *MATa ura3-Δ0 leu2-Δ0 trp1-Δ2 his3 lys2-128∂ tom1-Δ2(1873-2131)(+50, KanMX)* |
| 10233 | *tom1-intΔ4 (Δ1873-2022)* | *MATa ura3-Δ0 leu2-Δ0 trp1-Δ2 his3 lys2-128∂ tom1-Δ4(1873-2022)(+50 KanMX)* |
| 10234 | *tom1-intΔ5 (Δ1873-1972)* | *MATa ura3-Δ0 leu2-Δ0 trp1-Δ2 his3 lys2-128∂ tom1-Δ5(1873-1972)(+50 KanMX)* |
| 10270-4B | *tom1-intΔ9 (Δ1973-2131)* | *MATa ura3-Δ0 leu2-Δ0 trp1-Δ2 his3 lys2-128∂ tom1-Δ9(1973-2131)(+50 KanMX)* |
| 10291-7C | *tom1-intΔ10 (Δ1998-2059)* | *MATa ura3-Δ0 leu2-Δ0 trp1-Δ2 his3 lys2-128∂ tom1-Δ10 (1998-2059)(+50 KanMX)* |
| 10292-2C | *tom1-intΔ11 (Δ2031-2063)* | *MATa ura3-Δ0 leu2-Δ0 trp1-Δ2 his3 lys2-128∂ tom1-Δ11 (2031-2063)(+50 KanMX)* |
| Strains used for genomic analysis (RNA-seq, MNase-seq) | | |
| 8127-7-4 | *TOM1* | *MATa ura3-Δ0 leu2-Δ0 trp1-Δ2 his3 lys2-128∂* |
| 9843 | *tom1-intΔ1(1921-1947)* | *MATa ura3-Δ0 leu2-Δ0 trp1-Δ2 his3 lys2-128∂ tom1-Δ(1921-1947)(+50, KanMX)* |
| 9731-1-3 | *tom1-Δ0* | *MATa ura3-Δ0 leu2-Δ0 trp1-Δ2 his3 lys2-128∂ tom1-Δ0(::KanMX)* |
| Strains used for ChIP-seq (V5 immunoprecipitation) | | |
| 8127-7-4 | *TOM1 (untagged)* | *MATa ura3-Δ0 leu2-Δ0 trp1-Δ2 his3 lys2-128∂* |
| 10057 | *TOM1-12X-V5* | *MATa ura3-Δ0 leu2-Δ0 trp1-Δ2 his3 lys2-128∂ TOM1-12X-V5(KanMX)* |
| 10058 | *tom1-Δ1 (Δ1921-1947)-12XV5* | *MATa ura3-Δ0 leu2-Δ0 trp1-Δ2 his3 lys2-128∂ tom1-Δ(1921-1947)-12XV5(KanMX)* |
| Strains used for purifying different versions of Tom1p | | |
| 7382-3-4 | Protease-deficient *WT* | *MATa trp1 leu2 ura3 can1 pep4 prb1 his7* |
| 9736-K | Tom1p-PSc-PrtA | *MATa trp1 leu2 ura3 can1 pep4 prb1 his7 TOM1-PSc-PrtA(::KanMX)* |
| 9750 | Tom1p-PSc-PrtA | *MATa trp1 leu2 ura3 can1 pep4 prb1 his7 Gal1p-TOM1-PSc-PrtA(::KanMX)* |
| 10231 | Tom1p-ΔAD-PSc-PrtA | *MATa ura3-Δ0 leu2-Δ0 trp1-Δ2 his3 lys2-128∂ tom1-Δ2(1873-2131)-PSc-PrtA(::KanMX)* |
| 10232 | Tom1p-ΔAD-PSc-PrtA | *MATa trp1 leu2 ura3 can1 pep4 prb1 his7 Gal1p-tom1-Δ2(1873-2131)-PSc-PrtA(::KanMX)* |

| | | |
|---|---|---|
| 10285-12XHis | Tom1p-K1074-His12p | *MATa ura3-Δ0 leu2-Δ0 his3 lys2-128∂ tom1-K1074-Gly3-His12-D1075* |
| 10293-12XHis | Tom1p-K1074-His12 | *MATa trp1 leu2 ura3 can1 pep4 prb1 his7 Gal1p-tom1-K1074-His12-D1075* |

**Table S2. Oligonucleotides used for genomic manipulations and subcloning of fragments of Tom1p**

| | |
|---|---|
| To construct in-frame fusions to the C-terminus of the *TOM1* ORF, target fusion sequences were amplified with: | |
| MS 043 | GTTCACTATTATTGGCAATCAATGAAGGGCATGAAGGGTTTGGTCTTGCCCG ACGGATCCCCGGGTTAATTAAC |
| MS 044 | TCTAAAATACTTGGTTACATGGCGCTATAAATTTACACGAAAAATGATCATC ATCGATGAATTCGAGCTCGTTT |
| To transfer the *tom1-Δ0* allele from the standard collection ([Brachmann, 1998, 98144795]) to the A364a background, genomic DNA was amplified with: | |
| TF1 654 | CTCAACGCCGAAGGAATACCCTTC |
| TF1 655 | CCATTCGCTGTTTCAATATGGTTTTATAAGTGG |
| To integrate the pCORE cassette ([Storici, 2001 #50]) after *TOM1-S1943*, plasmid DS4664 was amplified with: | |
| TF1 714 | ATGCTATGGCATTTGTTGATAGTGATAACGGATTTGAAGTAGTATTTAGTGA GCTCGTTTTCGACACTGG |
| TF1 715 | TCTGAACGAGCATCATCTGCATCTTCTTCTCCCATATCATCATCTTCGTCTCC TTACCATTAAGTTGATC |
| To follow alleles of *TOM1* in crosses, markers were amplified with the following primers to integrate the PCR product 50 bp downstream of the *TOM1* ORF | |
| TF1 721 | TCATTTTTCGTGTAAATTTATAGCGCCATGTAACCAAGTATTTTAGAACGGG TTAATTAAGGCGCGCCAGATCTGTTTA |
| TF1 722 | CTTCCTTGGGCAAGTGTTGTATGGTTAAAGGTTTAAATAAAAGACGTTCTAT CATCGATGAATTCGAGCTCGTTTTCGAC |
| To integrate the pCORE cassette ([Storici, 2001 #50]) in the *TOM1* promoter, plasmid DS4664 was amplified with TF1658, TF1659, then the cassette was replaced with the *GAL1* promoter amplified with TF1661, TF1662: | |
| TF1 658 | GGTGAGAAAATAGTGCATATTTAGTTTACTTTTTGCCTTTGATTGAAAATAT ATATTCGAGCTCGTTTTCGACACTGG |
| TF1 659 | GCGGCGAGTTTCTCCTTTCTTGCCTTTTCACACCGAGTAAAAGCACCATTC CTTACCATTAAGTTGATC |
| TF1 661 | GGTGAGAAAATAGTGCATATTTAGTTTACTTTTTGCCTTTGATTGAAAATAT ATATTCTTATATTGAATTTTCAAAAATTCTTACTTTTTTTTTGGATGG |
| TF1 662 | GCGGCGAGTTTCTCCTTTCTTGCCTTTTCACACCGAGTAAAAAGCACCATTA TAGTTTTTTCTCCTTGACGTTAAAGTATAGAGG |
| To construct in-frame fusions to the N-terminus of the *TOM1* ORF, the promoter pCORE construct above was replaced with the product of PCR using: | |

| | |
|---|---|
| TF1960 | AGTGCATATTTAGTTTACTTTTTGCCTTTGATTGAAAATATATATTCATGCGACGGATCCCCGGGTTAATTAAC |
| TF1961 | CCAGCGGCGAGTTTCTCCTTTCTTGCCTTTTCACACCGAGTAAAAAGCACAAGTGGCGCGCCCGACTCGAGTGTAGAAT |
| To integrate the pCORE cassette ([Storici, 2001 #50]) after *TOM1-K1074,* plasmid DS4664 was amplified with TF2111, TF2112, then swapped for 12X-V5 or 12X His cassettes using TF2113, TF2114 or TF2118, TF2119, respectively. | |
| TF2111 | TTTATTTGCACATTATAAGGGATCTCTTTACGAGAATGACAAAAATAAAAGAGCTCGTTTTCGACACTGG |
| TF2112 | ATACCGTTGGACTCATCTATGTAGTTTATGTTATCAAGTGAGGATAAATCTCCTTACCATTAAGTTGATC |
| TF2113 | TTTATTTGCACATTATAAGGGATCTCTTTACGAGAATGACAAAAATAAAACGACGGATCCCCGGGTTAATTAAC |
| TF2114 | ATACCGTTGGACTCATCTATGTAGTTTATGTTATCAAGTGAGGATAAATCAAGTGGCGCGCCCGACTCGAGTGTAGAAT |
| TF2118 | TTTATTTGCACATTATAAGGGATCTCTTTACGAGAATGACAAAAATAAAATCTGGTGGAGGCGGTGGAGG |
| TF2119 | ATACCGTTGGACTCATCTATGTAGTTTATGTTATCAAGTGAGGATAAATCATGGTGATGATGATGGTGGTGATGA |

### Table S3. Cryo-EM data statistics

| **Data collection** | **Collection 1 (ΔAD)** | **Collection 2 (ΔAD)** | | **Collection 3 (FL)** | |
|---|---|---|---|---|---|
| Microscope | Krios G3 | Krios G3 | | Krios G3 | |
| Magnification | 81000X | 81000X | | 81000X | |
| Voltage (kV) | 300 | 300 | | 300 | |
| Electron exposure (e–/Å$^2$) | 47 | 47 | | 40 | |
| Defocus range (μm) | -0.8 to -2.0 | -0.8 to -2.0 | | -0.8 to -2.0 | |
| Pixel size (Å/pix) | 0.53 | 0.53 | | 0.55 | |
| **Data processing** | | | | | |
| Map description | **Tom1(ΔAD) (Closed)** | | **Tom1-FL (Closed)** | **Tom1-FL (Opening)** | **Tom1-FL (Opened)** |
| Micrographs used (no.) | 541 | 6536 | | 3,522 | |
| Particles picked (no.) | 136,510 | 1,160,670 | | 2,206,403 | |
| Particles retained (no.) | 462,711 | | 142,020 | 12,397 | 16,098 |
| **Map refinement** | | | | | |
| PDB/EMDB accession code | XXXX/XXX | | XXXX/XXX | N/A | N/A |
| Overall resolution (Å), FSC threshold 0.143 | 3.07 | | 3.97 | 7.79 | 7.10 |
| Map sharpening B-factor (Å2) | 117.3 | | 152.4 | 660.0 | 583.4 |
| **Model composition** | | | | | |
| Non-hydrogen atoms | 44632 | | 43300 | | |
| Protein residues | 2781 | | 2703 | | |
| **R.M.S. deviations** | | | | | |
| Bond lengths (Å) | 0.004 | | 0.004 | | |
| Bond angles (°) | 0.726 | | 0.943 (33) | | |
| **Validation** | | | | | |
| MolProbity score | 2.09 | | 2.61 | | |
| Clashscore | 11.49 | | 22.79 | | |
| Cβ outliers (%) | 0.00 | | 0.00 | | |
| Rotamer outliers (%) | 0.48 | | 0.83 | | |
| **Ramachandran plot** | | | | | |
| Favored (%) | 91.07 | | 77.87 | | |
| Allowed (%) | 7.78 | | 17.92 | | |
| Outliers (%) | 1.16 | | 4.2 | | |
| **Rama-Z (Plot Z-score, RMSD)** | | | | | |
| whole (N=2765) | -2.40 (0.16) | | -4.72 (0.15) | | |
| helix (N=1667) | -1.09 (0.12) | | -1.72 (0.13) | | |
| sheet (N=25) | -2.78 (1.08) | | -5.58 (0.71) | | |
| loop (N=1073) | -2.27 (0.19) | | -3.84 (0.16) | | |
| **CCvolume/mask** | 0.68/0.68 | | 0.68/0.69 | | |
| **CCpeaks/box** | 0.58/0.68 | | 0.62/0.76 | | |

**Table S4. X-ray crystallography data statistics**

| | |
|---|---|
| PDB ID | |
| Wavelength | 0.97946Å |
| Resolution range (outer shell) | 18–3.3 (3.6–3.3) |
| Space group | F 4(1) 3 2 |
| Unit cell (length;a=b=c) | 299.0 |
| Total reflections | 571306 (115734) |
| Unique reflections | 17852 (3583) |
| Multiplicity | 32.0 (32.3) |
| Completeness (%) | 99.9 (99.6) |
| Mean I/sigma(I) | 9.3 (1.4) |
| Wilson B-factor | 93.9 |
| R-merge | 0.58 (4.8) |
| R-meas | 0.59 (4.8) |
| R-pim | 0.104 (0.85) |
| CC1/2 | 0.995 (0.6) |
| CC* | 1 (0.96) |
| Reflections used in refinement | 17657 (2983) |
| Reflections used for R-free | 882 (142) |
| R-work | 0.228 (0.355) |
| R-free | 0.258 (0.375) |
| ligands | 15 |
| Solvent (atoms) | 4 |
| Protein residues | 407 |
| RMS(bonds) | 0.002 |
| RMS(angles) | 0.47 |
| Ramachandran favored (%) | 93.3 |
| Ramachandran allowed (%) | 6.45 |
| Ramachandran outliers (%) | 0.25 |
| Rotamer outliers (%) | 0.0 |
| Clashscore | 6.77 |
| Average B-factor | 84.10 |
| macromolecules | 83.98 |
| ligands | 117.9 |
| solvent | 53.55 |
